# Strain Level Microbial Detection and Quantification with Applications to Single Cell Metagenomics

**DOI:** 10.1101/2020.06.12.149245

**Authors:** Kaiyuan Zhu, Welles Robinson, Alejandro A. Schäffer, Junyan Xu, Eytan Ruppin, A. Funda Ergun, Yuzhen Ye, S. Cenk Sahinalp

## Abstract

The identification and quantification of microbial abundance at the species or strain level from sequencing data is crucial for our understanding of human health and disease. Existing approaches for microbial abundance estimation either use accurate but computationally expensive alignment-based approaches for species-level estimation or less accurate but computationally fast alignment-free approaches that fail to classify many reads accurately at the species or strain-level.

Here we introduce `CAMMiQ`, a novel combinatorial solution to the microbial identification and abundance estimation problem, which performs better than the best used tools on simulated and real datasets with respect to the number of correctly classified reads (i.e., specificity) by an order of magnitude and resolves possible mixtures of similar genomes.

As we demonstrate, `CAMMiQ` can better distinguish between single cells deliberately infected with distinct *Salmonella* strains and sequenced using scRNA-seq reads than alternative approaches. We also demonstrate that `CAMMiQ` is also more accurate than the best used approaches on a variety of synthetic genomic read data involving some of the most challenging bacterial genomes derived from NCBI RefSeq database; it can distinguish not only distinct species but also closely related strains of bacteria.

The key methodological innovation of `CAMMiQ` is its use of arbitrary length, doubly-unique substrings, i.e. substrings that appear in (exactly) two genomes in the input database, instead of fixed-length, unique substrings. To resolve the ambiguity in the genomic origin of doubly-unique substrings, `CAMMiQ` employs a combinatorial optimization formulation, which can be solved surprisingly quickly. `CAMMiQ`’s index consists of a sparsified subset of the shortest unique and doubly-unique substrings of each genome in the database, within a user specified length range and as such it is fairly compact. In short, `CAMMiQ` offers more accurate genomic identification and abundance estimation than the best used alternatives while using similar computational resources.

**Availability:** https://github.com/algo-cancer/CAMMiQ

## 1 Introduction

Advances in high throughput sequencing (HTS) have made it possible to generate millions of short reads in a few hours. An increased appreciation for the importance of microbes in human health and disease has prompted the generation of many metagenomic HTS datasets. For example, whole genome shotgun sequencing of the gut microbiome by the Human Microbiome Project provided important insight into the function and diversity of the human gut microbiome [1]. Similarly, the increase in HTS of human tissues represents another enormous source of metagenomic data because many of these human HTS datasets include reads from tissue-resident microbes, which have been shown to play an important role in many aspects of human disease, including tumorogenesis and the tumor response to therapy [2, 3, 4, 5, 6, 7].

Early approaches for analyzing metagenomic sequencing data were alignment based; reads were primarily searched in GenBank [8] through BLASTN [9] or custom built aligners such as GATK PathSeq (which, in return, is based on popular alignment tools) [10]. Unfortunately, the recent growth of HTS data and reference databases has made read search and alignment using BLASTN or PathSeq computationally infeasible. For example, a recent study which reported that microbial reads from tumors sequenced by The Cancer Genome Atlas (TCGA) can be used to build a classifier for cancer type [11] needed an “alignment-free” approach (Kraken [12]) due to the large number of samples analyzed. Kraken and other recent computational methods aim to identify and quantify specific genus in a metagenomic sample faster through a number of algorithmic techniques are summarized in [13]. Even though these methods address the problem of computational scalability, available alignment-free 1tools do not match the accuracy of alignment-based tools. For example, another recent paper on microbial reads from single cell RNA-seq (scRNA-seq) datasets to distinguish cell type specific intracellular microbes from extracellular and contaminating microbes [14] had to use PathSeq. This was because of the relatively small number of microbial reads per cell, to classify as many reads as possible at the genus and species levels - the resolution of available alignment-free methods were too limited for this task. The distinct approaches taken by these two studies illustrate the trade-offs inherent in the alignment-based and alignment-free methods to metagenomic identification and quantification.

One additional way to achieve speedup is through the reduction of the size of the sequence database used. Methods in this direction may align reads only to *marker genes,* a relatively small collection of clade-specific, single-copy genes, instead of the full reference genomes [15, 16], through the use of available read mapping techniques. Such methods need to obtain the collection of marker genes by employing information beyond what is offered by the database itself - which is not always possible. Furthermore, since these methods identify and use only a handful of marker genes on each genome, many of the reads in the HTS data can not be utilized, implying low specificity. As a consequence, species with low abundance within the sample may be difficult to identify (let alone correctly quantify) because variation in HTS coverage may result in a few or no reads originating from the marker genes.

Alignment-free methods have been available in the context of string comparison for a long time [17, 18, 19] and they have found applications in bioinformatics workflows before the emergence of HTS [20, 21]. Applications of alignment-free methods to metagenomic HTS data typically rely on k-mer “matches” to return a taxonomic assignment for every read. These applications either assign a read to the lowest taxonomic rank possible (determined by the specificity of the read’s *k*-mers) [22, 12, 23, 24], or to a pre-determined genus, species, or strain taxonomic level [25, 26]. In contrast to marker gene based methods, *k*-mer based applications can usually assign each given read to the correct genus. As the value of *k* becomes larger, more reads can be assigned a unique label at the desired taxonomic rank, however, with growing *k*, the space requirement of these methods grows very quickly - which, in the worst case, can imply a factor *k* increase on the space requirements of the original database. The large memory footprint to maintain the entire *k*-mer profile of each species can be reduced through hashing or subsampling the *k*-mers [27, 28]; however this would result in loss of accuracy. In addition to methods based on exact *k*-mer counts, it is also possible to assign metagenomic reads to bacterial genomes by employing species-specific sequence features (e.g. short *k*-mer distribution or GC content) [29, 30, 31, 32, 33], although methods that employ this approach are typically not very accurate at species level or (especially) at strain level assignment.

The ability to identify and quantify distinct pathogenic strains in a microbial sample has many applications in diagnosing and treating infectious diseases [34]. In particular, mixed infections that are caused by multiple strains of a bacterial species [35, 36], have been described for tens of bacterial species [37]. It is estimated that a high fraction of *M. tuberculosis* and *Staphylococcus aureus* patients are infected with multiple strains [34], each with distinct different drug susceptibility and antibiotic resistance profiles. Clearly accurate analysis of microbial sequence data at the strain level would be highly helpful for identifying mixed infections. Additionally, strain-level analysis can be important for distinguishing pathogenic strains from non-pathogenic strains [38] and for tracking food-borne pathogens [39].

As have been summarized above, neither the alignment free, k-mer based methods, nor the marker-gene based tools take into consideration the distribution of the reads over the genome of the species they are assigned. In fact, the alignment-free methods do not even take into account the length of the genomes, but are based on (unnormalized) read counts. In reality, provided that the sequence data to be analyzed is genomic, the distribution of reads generated by high throughput sequencing from a given species or strain should be roughly uniform. This principle is fundamental to a number of *isoform abundance estimation* methods, which aim to solve a very similar problem [40, 41, 42]. Interestingly, this constraint is underutilized in the context of metagenomic analysis. One exception is the network flow based approach utilized by, e.g. [43], which (implicitly) establishes a reference guided assembly of the reads into the genomes of the species involved. As such it is quite accurate but is very slow. Another method in this direction considers the uniformity of coverage across k-mers with each genome to reduce false positive calls [44], which improves the running time at a moderate loss of accuracy.

In contrast to the metagenomic species identification and quantification methods summarized above, there are also tools to determine the likely presence of a long genomic sequence (e.g. the complete or partial genome of a bacterial species) in a given metagenomic sample [45, 46, 47, 48]. Even though these tools solve an entirely different problem, methodologically they are similar to the *k*-mer based species identification and quantification tools such as [12, 26] in the sense that they build a succinct index on the database (this time comprised of the the metagenomic read collection) and query this index without explicit alignment. And because of their design parameters, these tools can not perform abundance estimation for a given species.

### Our Contributions

In this paper we describe `CAMMiQ` (Combinatorial Algorithms for Metagenomic Microbial Quantification), a new computational method to maintain/manage a collection of *m* (bacterial) genomes 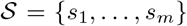, each assembled into one or more strings/contigs, representing a species, a particular strain of a species, or any other taxonomic rank. `CAMMiQ`’s data structure can answer queries of the following form: given a query set 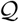 of HTS reads obtained from a mixture of genomes or transcriptomes, each from 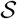, identify the genomes in 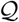, and, in case the reads are genomic, compute their relative abundances. Our data structure is not only very efficient in terms of its querying time, but is also shown to be very accurate, through simulations. The key novel feature of our data structure is its utilization of substrings that are present in at most *c* genomes (*c* > 1) in 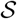. There are alignment free methods that utilize unique *k*-mers in genomes for metagenomic analysis [12, 26, 44] already. However, our data structure is the first to consider those substrings that are present in two or possibly more genomes, for increasing the proportion of reads it can utilize and thus improving sensitivity. Because of this novel feature, our data structure can accurately identify genomes at subspecies/strain level.

In order to assign each read in 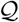 that includes an “almost-unique” substring (i.e. present in at most *c* genomes) to a genome, our data structure solves an integer linear program (ILP) - that simultaneously infers which genomes are present in 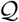 and, (if the reads are genomic) their relative abundances. Specifically, the objective of the ILP is to identify a set of genomes, in each of which the coverage of the almost-unique genomic substrings is (approximately) uniform.

One novel feature of our data structure is its use of *shortest* substrings (present in at most *c* genomes) - rather than fixed length “*k*-mers”.^1^ This feature also increases the number of reads utilized by our data structure since some reads may include no almost-unique *k*-mer but instead longer substrings that are almost-unique. It also reduces the number of substrings and thus features (not to be confused with the number of reads) to be handled by the ILP: this is because, rather than considering multiple overlapping (almost-)unique *k*-mers, our data structure uses a single shorter substring shared by all of them. The ILP formulation further reduces the number of almost-unique substrings it considers by maintaining a maximally sparsified set of substrings that guarantee any potential read from a genome in 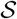 that is almost-unique includes one such substring. In the remainder of the paper, we set the value of *c* to 2, so as to consider not only unique but also “doubly-unique” substrings of the genomes in 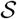. Even though this choice turned out to be sufficiently powerful for the datasets we experimented with, it is easy to generalize our approach to *c* > 2 (i.e. to triply-unique, etc. substrings).

As a final contribution, we provide sufficient conditions to identify and quantify genomes in a query correctly, through the use of unique substrings/*k*-mers, provided the reads are error-free. Although this is a purely theoretical result, to the best of our knowledge it is the first of its kind for metagenomic data analysis, and is valid for `CAMMiQ` for the case *c* =1 and other unique substring based methods such as CLARK and KrakenUniq. `CAMMiQ`’s use with *c* = 2 is primarily advised for cases where these conditions are not satisfied.

We show that `CAMMiQ` is not only faster but also more accurate than GATK PathSeq on scRNA-seq data obtained from monocyte-derived dendritic cells (moDCs) infected with distinct *Salmonella* strains analyzed in [14]. Then we demonstrate the comparative advantage of `CAMMiQ` against some of the best used metagenomic tools on synthetic genomic HTS data from “challenging” microbial strains we derived from the NCBI RefSeq database.

## 2 Algorithmic Formulation

The input to `CAMMiQ` is a set of *m* genomes 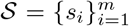, not necessarily all from the same taxonomic level (each genome here may be associated with a genus, species, subspecies or strain) to be indexed. Although we describe `CAMMiQ` for the case where each 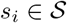 is a single string, we do not assume that the genomes are fully assembled into a single contig; rather the string representing a genome could simply be a concatenation of all contigs from species *i* and their reverse complements with a special symbol $_*i*_ between each contig. We call 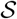 the input *database* and *i* ∈ {1, ···, *m*} the *genome ID* of string *s*_*i*_.

A query for `CAMMiQ` involves a set of reads 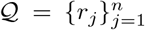 representing a metagenomic mixture. For simplicity we describe `CAMMiQ` for reads of length *L*, however our data structure can handle reads of varying length. Given 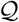, the goal of `CAMMiQ` is to identify a set of genomes 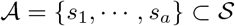 and their respective abundances *p*_1_, ···, *p*_*a*_ which “best explain” 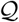. This is achieved by assigning (select) reads *r*_*j*_ to genomes *s*_*i*_ such that the implied coverage of each genome 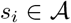 is uniform across *s*_*i*_, with *p*_*i*_ as the mean.

In its simplest form, `CAMMiQ` builds a succinct index for input database 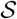, so as to handle queries in the following form. Given 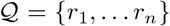 find the smallest set of genomes 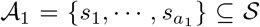, such that the set of *unique L*-mers in 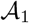 includes all reads in 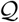 that are identical to a unique *L*-mer in 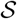. For any 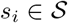, call an *L*-mer in *s*_*i*_ unique in 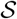 if it does not appear in any other *s*_*i′*_. We denote by *U*_*i,L*_, the set of all unique L-mers in *s*_*i*_. Then 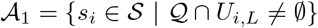.

The above query type may not be powerful enough to identify in 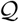 those genomes in 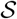 that are very similar (with respect to sequence composition) to other genomes and thus do not include many unique *L*-mers. Such a genome *s*_*i*_ would be especially problematic if it is low in abundance, since the chances of 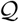 to include any unique *L*-mer from *U*_*i,L*_ will be low. As a solution to this problem, `CAMMiQ` also features a second type of query, which is more general since it involves both unique and *doubly-unique L*-mers of genomes *s*_*i*_. We call an *L*-mer doubly-unique in 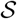 if it appears in exactly two genomes *s*_*i*_ and *s*_*i′*_ and denote the set of doubly-unique *L*-mers in *s*_*i*_ by *D*_*i,L*_. Given query set 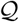, `CAMMiQ`’s more general query asks to compute 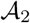, the smallest subset of 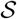 which include all reads in 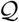 identical to unique and doubly-unique *L*-mers in 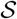. Note that, necessarily 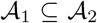. Also note that, by involving doubly-unique *L*-mers, this more general query may have a chance to capture those genomes in 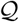 which have low abundance and have a few unique *L*-mers.

Unfortunately, the index structure necessary to support the second type of queries is larger than that for the first type of queries. Furthermore, this query type could still produce inaccurate results in the presence of read errors. For handling read errors (at noise rates commonly observed in Illumina data) `CAMMiQ` finally features a third type of query, which is even more general since it involves the “shortest” unique and doubly-unique substrings of reads in 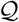. We call a substring of a genome 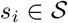 a shortest unique substring, if it is unique to *s*_*i*_, has length in the range [*L*_*min*_, *L*_*max*_], and has no substring that is a shortest unique substring; this definition can be extended to shortest doubly-unique substrings as well. This query thus asks to compute the smallest subset 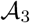 of 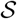 which include these substrings, with the constraint that the “coverage” of these substrings in each genome 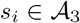 is “uniform”. The query also asks to compute the relative abundance of each genome in 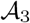 as will be described later.

By considering shorter unique substrings, our most general query has a higher chance of observing them within 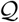 (since a substring of length *L′* < *L* is included in *L* − *L′* + 1 *L*-mers in the genome). This query may also imply that a smaller fraction of unique or doubly-unique substrings in 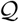 would be subject to read errors (since it is likely that each read may include more than one such unique substring and it is likely that at least one such unique substring will be error free). Furthermore the index structure necessary to maintain the entire set of shortest unique and doubly-unique substrings of genomes in 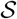 is fairly large. As a result of this, we build the index on a maximally sparsified set of shortest unique and doubly-unique substrings of each 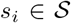 which ensure that each read in 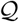 that can be attributed to at most two genomes includes at least one such substring (see the Supplementary Methods for more details).

An index on this sparsified set of shortest unique and doubly-unique substrings is sufficiently powerful for `CAMMiQ` to answer all three types of queries, i.e. it can efficiently compute the sets 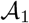, 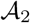 and 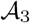. For all three query types `CAMMiQ` first identifies for each read *r*_*j*_ all unique and doubly-unique substrings it includes; it then assigns *r*_*j*_ to the one or two genomes from which these substrings can originate^2^. To compute 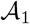 `CAMMiQ` can simply return the collection of genomes which receive at least one read assignment. To compute 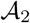 `CAMMiQ` needs to solve the “set cover” problem, or more precisely, its dual, the “hitting set” problem where genomes form sets and indexed strings that appear in query reads form the items to be covered. To compute 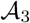 `CAMMiQ` solves the combinatorial optimization problem that asks to minimize the variance among the number of reads assigned to each indexed substring of each genome - the solution indicates the set of genomes in 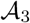 along with their respective abundances.

Details on the composition as well as the construction process for `CAMMiQ`’s index are discussed in Section 2.1. The query processing of `CAMMiQ`, which proceeds in two stages are discussed in Sections 2.2 and 2.3: The first stage assigns reads to specific genomes - which is sufficient for computing sets 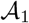 and 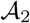. See Section 2.2 for the criteria we use for assigning a read to a genome, based on the indexed substrings it includes. The second stage introduces the combinatorial optimization formulation to compute 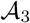 as a response to the most general query type. See Section 2.3 for details.

### 2.1 Index Construction

In order to respond to the three types of queries described above, we preprocess unique and doubly-unique substrings of genomes in 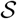 to form an index structure as follows. Let 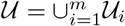 and 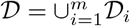 where 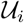 and 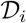 are the substrings from genome *s*_*i*_ whose lengths are within the range than [*L*_min_, *L*_max_ ≤ *L*], such that each 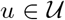 is present in at most one genome and each 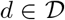 is present in at most two genomes in 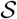. See below for a detailed definition for the uniqueness of a substring. If our goal is to compute 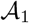 or 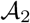 only (i.e. respond to the first and second type of queries), we can simply set *L*_min_ = 0, *L*_max_ = *L* and maintain the corresponding substring collections 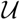 and 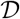 in our index. For computing 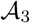 (i.e. responding to the third type of queries - in addition to the first and second types), we actually use the length constrained definitions of substrings for computing 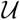 and 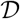. As mentioned earlier, we then sparsify 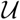 and 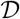 as much as possible by maintaining only one representative substring among those that are in close proximity within a genome, and discarding the rest (see “Subsampling unique substrings” below for details). We start with formal definitions and some notation.

#### Notation and definitions

Let *s* = *s*_1_$_1_ ∘ ··· ∘ *s*_*m*_$_*m*_ denote the string obtained by concatenating the input reference genomes 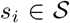 and let *M* = |*s*| = ∑_*i*_ |*s*_*i*_| denote its length. A substring of *s* is a string in the form *s*[*l* : *r*] = *s*[*l*]*s*[*l*+1] ··· *s*[*r*]. With a slight abuse of notation we denote by *s*_*i*_[*l* : *r*] not the actual substring of *s*_*i*_ including its *l*^*th*^ to *r*^*th*^ symbols, but rather the substring of *s* including its *l*^*th*^ to *r*^*th*^ symbols, with the provision that all these symbols are within the representation of *s*_*i*_ in *s*. We denote by an ℓ-mer a string of length ℓ. The suffix of s that starts at position *i* is denoted `suf`[*i*] = *s*[*i*, ···, *M*]. In what follows, we use the *generalized enhanced suffix array* of *s* which is composed of three parts. (i) The *suffix array* `SA` of *s*, which is comprised of the positions 1, 2, ···, *M*, sorted in increasing lexicographical order of the corresponding suffixes `suf`[*i*], *i* = 1, 2, ···, *M*. That is, `SA`[*i*] = *j* indicates that `suf`[*j*] is the *i*-th smallest suffix in lexicographical order. In addition, we denote `SA`^−1^[*j*] = *i* if `SA`[*i*] = *j*. (ii) The *longest common prefix array,* `LCP`, contains in its *i*-th position the length of the longest common prefix of `suf`[`SA`[*i*]] and `suf`[`SA`[*i*−1]], for 2 ≤ *i* ≤ *M* (and `LCP`[1] = 0). (iii) Finally, the *generalized suffix array* `GSA` contains the genome ID of each suffix `suf`[`SA`[*i*]]. All of the above arrays can be constructed in linear time: the first data structure that can be constructed in linear time, with the ability to determine whether a given substring is unique to a “document” (i.e. a genome in our context) in a collection of documents, and compute the shortest unique substring of a document that ends in a particular position, in time proportional to the substring length is the augmented suffix tree of Matias et al. [51]. Once an augmented suffix tree is computed, it can be trivially reduced to the above described enhanced suffix array in *O*(*M*) time - which can also be constructed without the use of suffix trees to achieve a constant factor improvement in memory [52, 53].

We denote by *u*_*i*_(*l*, *r*) the substring *s*_*i*_[*l* : *r*] that is *unique* to genome *s*_*i*_; formally, this indicates that there exists no substring *u*_*j*_(*l′*, *r′*) on any genome *s*_*j*_ ≠ *s*_*i*_ such that *u*_*i*_(*l*, *r*) = *u*_*j*_(*l′*, *r′*). We call *u*_*i*_(*l*, *r*) a shortest unconstrained unique substring, if none of its substrings are unique. Similarly, we denote by *d*_*i*_(*l*, *r*) the substring *s*_*i*_ [*l* : *r*] that is *doubly-unique* to genome *s*_*i*_ and one other genome, say *s*_*j*_; formally, this indicates that there is exactly one genome *s*_*j*_ which includes the substring *d*_*j*_(*l′*,*r′*), i.e., *s*_*i*_[*l* : *r*] = *s*_*j*_[*l′* : *r′*] for some *l′*,*r′*. Clearly, any superstring of a unique substring is still unique and any superstring of a doubly-unique substring is either unique or doubly-unique. We call *d*_*i*_(*l*, *r*) a shortest unconstrained doubly-unique substring of *s*_*i*_ and some other genome *s*_*j*_, if none of its substrings are doubly-unique.

For our purposes, we need to constrain the shortest unique and doubly-unique substrings with length upper bound *L*_max_ and lower bound *L*_min_. Under these constraints, we call any shortest unconstrained unique substring *u*_*i*_(*l*, *r*) a shortest unique substring if *L*_max_ ≥ *r* − *l* + 1 > *L*_min_. We also call a unique substring *u*_*i*_(*l*, *r*) a shortest unique substring if *r* − *l* + 1 = *L*_min_. Similarly, we call any shortest unconstrained doubly-unique substring *d*_*i*_(*l*, *r*) a shortest doubly-unique substring if *L*_max_ ≥ *r* − *l* + 1 > *L*_min_. Again, we call a doubly-unique substring *u*_*i*_(*l*, *r*) a shortest doubly-unique substring if *r* − *l* + 1 = *L*_min_ as well. We say an *L*-mer *s*_*i*_[*l* : *l* + *L* − 1] includes a unique substring *s*_*i*_[*l′* : *r′*], or, conversely, a unique substring *s*_*i*_[*l′* : *r′*] covers an *L*-mer *s*_*i*_[*l* : *l* + *L* − 1] if *l′* ≥ *l* and *r′* ≤ *l* + *L* − 1. As such, we call an *L*-mer unique if it includes a unique substring. We can generalize these definitions to the notion of an *L*-mer including a doubly-unique substring, or conversely, a doubly-unique substring covering an *L*-mer, and thus making the *L*-mer itself doubly-unique - provided that it is not unique.

#### Algorithmic framework to compute shortest unique substrings

It is quite simple to compute the shortest unique and doubly-unique substrings in 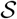 in *O(M)* time by using the augmented suffix tree described in [51]. A similar running time can also be achieved through the use of a suffix array, as discussed by [54] for a single document (i.e. genome). We slightly generalize this to handle multiple genomes as follows. The key observation we use is that given a position *l*, the shortest unique or doubly-unique substring of *s*_*i*_ that starts at *l* (i.e. *u*_*i*_(*l*,*r*) or *d*_*i*_(*l*, *r*)) is the shortest unique, or respectively doubly-unique prefix of `suf`[*l*]. In this way the problem can be reduced to searching for the longest common prefix of `suf`[*l*] with any other suffix from another genome (i.e., any genome with ID ≠ `GSA`[`SA`^−1^[*l*]]) for each 1 ≤ *l* ≤ *M*. Let lcp(*x,y*) denote the longest common prefix of two suffices *x* and *y*; then we define:

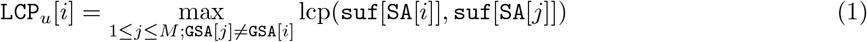

and

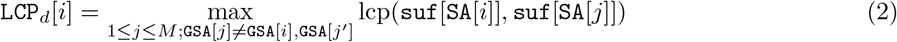

where *j′* indicates a suffix `suf`[*j′*] with `GSA`[*j′*] ≠ `GSA`[*i*], which maximizes lcp(`suf`[`SA`[*i*]], `suf`[`SA`[*j′*]]); note that any value of *j′* that maximizes lcp(`suf`[`SA`[*i*]], `suf`[`SA`[*j′*]]) will imply the same value for `LCP`_*d*_[*i*]. `CAMMiQ` maintains an array `SU` (of length *M*) such that `SU`[*r*] = *l* if *u*_*i*_(*l, r*) is a shortest unique substring. In order to compute `SU`, each of its entries is initially set to 0 and for each *i* = 1, … *M*, one entry of SU is updated as

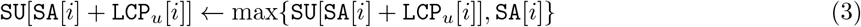

Similarly, `CAMMiQ` maintains an array `SD` (again of length *M*) such that `SD`[*r*] = *l* if *d*_*i*_(*l,r*) is a shortest doubly-unique substring. Again each entry of `SD` is initially set to 0 and then for each *i* = 1,… *M*, one entry of `SD` is updated as

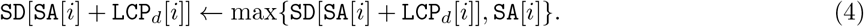

See Supplementary Section 5.1 for further details.

#### Computing LCP_*u*_ and LCP_*d*_

Given `GSA`[*i*_1_, ···,*i*_2_] a subarray of `GSA`, let *d*_GSA_ (*i*_1_,*i*_2_) be the number of distinct genomes the entries in this subarray belong to, i.e. *d*_GsA_(*i*_1_,*i*_2_) = |{`GSA`[*i*_1_], ···, `GSA`[*i*_2_]}|. We can now compute `LCP`_*u*_ and `LCP`_*d*_ in linear time as follows.

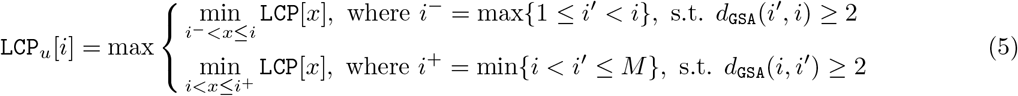

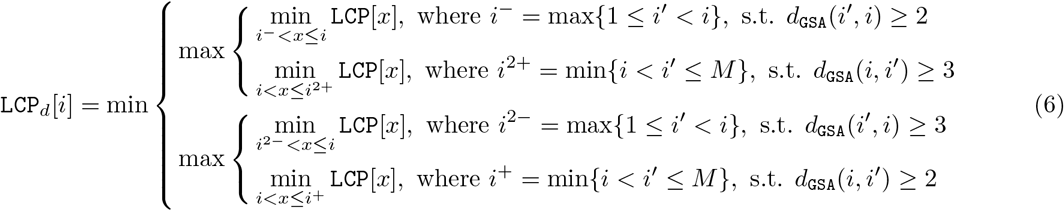

Note that to introduce a minimum length constraint *L*_min_ on unique and doubly-unique substrings, each `LCP`_*u*_[*i*] is (re)set to max{*L*_min_ − 1, `LCP`_*u*_[*i*]} and respectively each `LCP`_*d*_[*i*] is (re)set to max{*L*_min_ − 1, `LCP`_*d*_[*i*]}. Then, to ensure that each shortest doubly-unique substring occurs in exactly two genomes (and not one), we set `LCP`_*d*_[*i*] = ∞ to in case the above procedure ends up with `LCP`_*d*_[*i*] = `LCP`_*u*_[*i*]. See Supplementary Section 5.2 for the proof of correctness and a running time analysis for the computation of `LCP`_*u*_ and `LCP`_*d*_.

#### Subsampling unique substrings

As defined above, the array SU maintains the indices of all shortest unique substrings; some of these substrings may have large overlaps with others and thus are redundant in assessing the uniqueness of a read in the query. Let 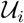 be the collection of all unique substrings on genome *s*_*i*_. Then, in order to reduce the index size, `CAMMiQ` aims to compute a subset 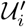 of 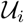, consisting of the minimum number of shortest unique substrings that cover every unique *L*-mer on *s*_*i*_. `CAMMiQ` also aims to compute a subset 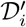 of 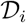, consisting of the minimum number of shortest doubly-unique substrings that cover every doubly-unique *L*-mer on *s*_*i*_. This is all done by greedily maintaining only the rightmost shortest unique or doubly-unique substring in any *L*-mer of a genome in 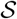. In the remainder of the paper we denote the number of unique substrings in subset 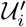 by 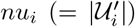 and respectively the number of doubly-unique substrings in subset 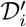 by 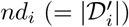; we denote the number of unique *L*-mers on *s*_*i*_ by 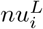 and respectively the number of doubly-unique *L*-mers on *s*_*i*_ by 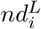. As we prove in Supplementary Section 5.3, this greedy strategy we employ can indeed obtain the minimum number of shortest unique substrings to cover each unique *L*-mer, provided that each substring in 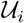 occurs only once *s*_*i*_.

#### Index Organization

Consider the set of unique substrings Let 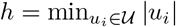 be the minimum length of all shortest unique substrings (*h* is automatically set to *L*_min_ if the minimum length constraint is imposed). `CAMMiQ` maintains a hash table which maps a distinct *h*-mer *w* to a bucket containing all unique substrings *u*_*i*_ which have *w* as a prefix. Within each bucket, the remaining suffices of all unique substrings *u*_*i*_, i.e. *u*_*i*_[*h* + 1 : |*u*_*i*_|], are maintained in a trie so that each leaf contains the corresponding genome ID. For each read *r*_*j*_ in the query, `CAMMiQ` considers each substring of length h and its reverse complement and computes its hash value (in time linear with *L* through Karp-Rabin fingerprinting [55]). If the substring has a match in the hash table, then `CAMMiQ` tries to extend the match until a matching unique substring is found, or finds no match. (Note that the processing for doubly-unique substrings is identical to that for unique substrings.) See Figure 1 for an overview of the index structure. Also see Section 2.2 below for the use of unique and doubly-unique substrings identified for each read to answer the query.

**Figure 1:**
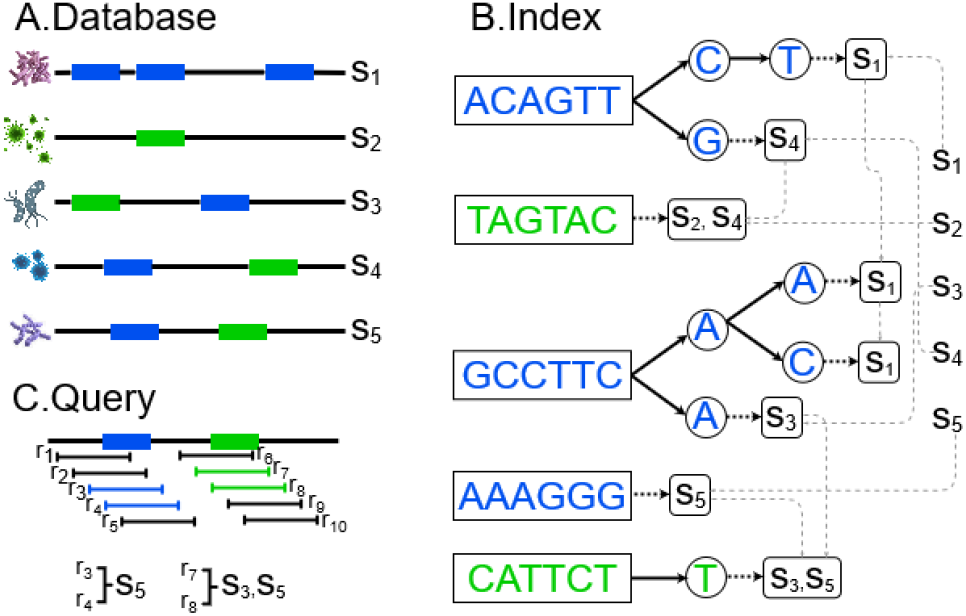
Overview of `CAMMiQ`’s index structure. Strings in blue are unique substrings and those in green are doubly-unique substrings.

### 2.2 Query Processing Stage 1: Preprocessing the Reads

Given the index structure on the sparsified set of shortest unique and doubly-unique substrings of genomes in 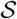, we handle each query 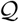 in two stages. In Stage 1, we preprocess each read 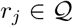 as follows. Consider the collection of genome sets associated with each unique and doubly-unique substring in the read *r*_*j*_. If the intersection of these sets is empty, we discard the read since these substrings are “conflicting” (see below for a short discussion on these conflicts). If the intersection is non-empty, for each unique substring *u*_*i*_ and each doubly-unique substring *d*_*i*_ in *r*_*j*_, we increase by 1 the associated counter *c*(*u*_*i*_) and respectively *c*(*d*_*i*_) we maintain. These counters indicate the number of “conflict-free” reads that include each such substring in 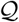. As described below, these counters will be essential to actual query processing in Stage 2.

1. Identify the set 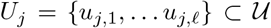 of unique substrings in *r*_*j*_; similarly identify the set 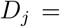 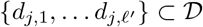 of doubly-unique substrings in *r*_*j*_.
2. a. If *U*_*j*_ = *D*_*j*_ = ∅ then discard *r*_*j*_.
b. If *U*_*j*_ ≠ ∅ but it includes a pair of unique substrings *u*_*j,i*_ and *u*_*j,i′*_ which originate from different genomes, then discard *r*_*j*_.
c. If *U*_*j*_ ≠ ∅ and all its unique substrings originate from the same genome *s*_*k*_, however *D*_*j*_ includes a substring *d*_*j,i*_ which can not originate from *s*_*k*_, then again discard *r*_*j*_.
d. If *U*_*j*_ = ∅ and the intersection of the set of genomes from where the substrings in *D*_*j*_ can originate is empty, then also discard *r*_*j*_.
e. If on the other hand,

i. *U*_*j*_ ≠ ∅, all its unique substrings originate from the same genome *s*_*k*_, and each doubly-unique substring *d*_*j,i′*_ ∈ *D*_*j*_ can originate from *s*_*k*_, or
ii. *U*_*j*_ = ∅, however the intersection between the genomes where the doubly-unique substrings of *r*_*j*_ can originate from is comprised of only one genome, *s*_*k*_, or
iii. *U*_*j*_ = ∅ and *d*_*j*_ is comprised of doubly-unique substrings that can only originate from the same pair of genomes *s*_*k*_ and *s*_*k′*_, then then increase *c*(*u*_*j,i*_) by 1 for each *u*_*j,i*_ ∈ *U*_*j*_ and *c*(*d*_*j,i*_) by 1 for each *d*_*j,i*_ ∈ *D*_*j*_.

The above counters are sufficient to compute the set 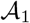 as well as 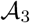, the answer to our most general query type. For computing 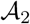, `CAMMiQ` additionally maintains a counter *d*(*s*_*k*_, *s*_*k′*_) for each pair of genomes *s*_*k*_, *s*_*k′*_, indicating the number of reads in 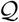 that can originate both from *s*_*k*_ and *s*_*k′*_; the value of this counter needs to be increased for each pair of involved genomes by 1 in case (iii) above.

The preprocessing stage described above eliminates those reads that include conflicting unique or doubly-unique substrings - conflicting in the sense that they are associated with different genomes. There are two main reasons for observing such conflicts: (i) sequencing errors, (ii) the presence of reads in the query from genomes that are not in 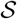. By eliminating these conflicting reads, we reduce the chances of mis-identifying the genomes from which they may originate.

In short, the read preprocessing stage produces two vectors 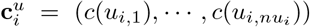 and 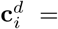 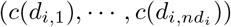 that indicate the number of (conflict-free) reads that include each unique and doubly-unique substring on each genome *s*_*i*_. One can use these vectors to compute 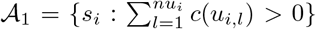. Additionally, through the use of the counter *d*(*s*_*k*_, *s*_*k′*_) maintained for each pair of genomes *s*_*k*_, *s*_*k′*_ one can compute 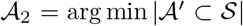 such that (i) 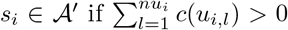 and (ii) 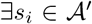, if *d*(*s*_*k*_, *s*_*k′*_) > 0 then either *i* = *k* or *i* = *k′*. This is basically the solution to the hitting set problem we mentioned earlier, whose formulation as an integer linear program (ILP) is well known [56]. From this point on, our main focus will be computing 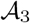 (as well as the relative abundances of the genomes in 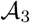) for which we introduce an ILP formulation described below.

### 2.3 Query Processing Stage 2: ILP Formulation

`CAMMiQ` computes the list of genomes in the query as well as their abundances through an integer linear program (ILP) described below. Let *δ*_*i*_ = 0/1 be the indicator for the absence or presence of the genome *s*_*i*_ in 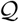. The ILP formulation assigns a value to each *δ*_*i*_ and also computes for each *s*_*i*_ its abundance *p*_*i*_ in the range [*p*_min_,*p*_max_].

**Minimize**

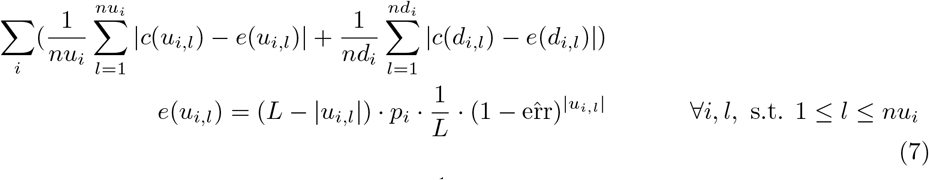

**Subject to**

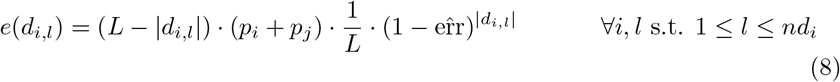

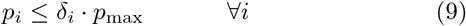

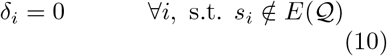

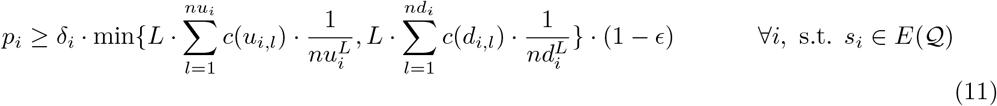

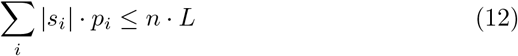

The objective of the ILP is to minimize the sum of absolute difference between the expected and the actual number of reads to cover a unique or doubly-unique substring. Since each genome may have different number of unique and doubly-unique substrings, this difference is normalized w.r.t. *nu*_*i*_ or *nd*_*i*_. Constraints (7) and (8) define the expected number of reads to cover a particular unique substring *u*_*i,l*_ or doubly-unique substring *d*_*i,l*_ - here *p*_*j*_ is the abundance of genome *s*_*j*_ which also includes *d*_*i,l*_. Here we denote by 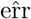 the estimated read error (specifically substitution) rate per nucleotide, and denote by |*w*| the length of a substring *w*. Constraint (9) ensures that the abundance *p*_*i*_ of a genome is 0 if *δ*_*i*_ = 0. Constraint (10) ensures that the solution to the above ILP excludes those genomes whose counters for unique and doubly-unique substrings add up to a value below a threshold - so as to reduce the size of the solution space. More specifically, given a threshold value *α*, the constraint excludes those genomes *s*_*i*_ that are not in the set of genomes 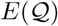 whose counters for its unique substrings add up to a value above a 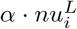, and doubly unique substrings add up to a value above a 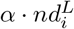. More formally, 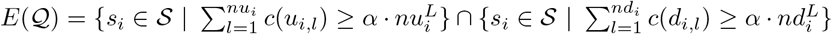. Constraint (11) enforces a lower bound on the coverage (and thus the abundance) of each genome *s*_*i*_ in the solution to the above ILP (namely, with *δ*_*i*_ = 1), which must match the coverage 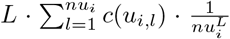 and 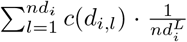 resulting from the number of reads in 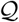 that include a unique and doubly-unique substring respectively, i.e., it must be at least (1 − *ϵ*) times the smaller one above for a user defined *ϵ*. Constraint (12) enforces an upper bound on the coverage (and thus the abundance) of each genome *s*_*i*_ in the solution to the above ILP, through making the sum (over each *s*_*i*_) of the number of reads produced on *s*_*i*_ based on *p*_*i*_ not exceed the total number of reads *n*. Collectively, the last two constraints ensure that the abundance *p*_*i*_ computed from the ILP matches what is (i.e., the coverage based on read counts) given by 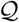. Note that the absolute values in the objective can be removed by introducing a new variable *γ*(*u*_*i,l*_) ≥ max{*c*(*u*_*i,l*_) − *e*(*u*_*i,l*_), *e*(*u*_*i,l*_) − *c*(*u*_*i,l*_)}.

### 2.4 When to Use Unique Substrings - the Error Free Case

We now provide a set of sufficient conditions to guarantee the (approximate) performance that can be obtained (with high probability) in metagenomic identification and quantification by the use of unique substrings only. These conditions apply to `CAMMiQ` when *c* = 1, as well as CLARK, KrakenUniq and other similar approaches. In case these conditions are not met, it is advisable to use `CAMMiQ` with *c* ≥ 2.

Suppose that we are given a query 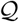 composed of n error-free reads of length *L*, sampled independently and uniformly at random from a collection of genomes 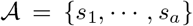 according to their abundances *p*_*1*_, ···, *p*_*a*_. More specifically, suppose that our goal is to answer query 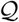 by computing 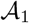, along with an estimate for the abundance value *p*_*i*_ for each 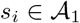, calculated as the weighted number of reads assigned to *s*_*i*_ according to the procedure described in Section 2.2. Then, the L1 distance between the true abundance values and this estimate will not exceed a value determined by *n* (number of reads), *a*, and *q*_min_, the minimum (normalized) proportion of unique *L*-mers among these genomes. For a given failure probability *δ* and an upper bound on L1 distance *ϵ*, this translates into sufficient conditions on the values of *n*, *a* and *q*_min_ to ensure acceptable performance by the computational method in use.

#### Theorem 1.

*Let* 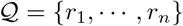 *be a set of n error-free reads of length L, each sampled independently and uniformly at random from all positions on a genome* 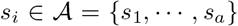 *where s*_1_, ···, *s_a_ is distributed according to their abundances p*_1_, ···, *p_a_* > 0. *Let* 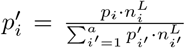 *be the corresponding “unnormalized” abundance of p_i_ for i* = 1, ···, *a, where* 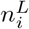 *denotes the total number of L-mers on s_i_. Let q*_1_, ···, *q_a_* > 0 *be the proportion of unique L-mers on s*_1_, ···, *s_a_ respectively; p*_min_ = min{*p*_1_, ···, *p_a_*}; *q*_min_ = min{*q*_1_, ···, *q_a_*}. *Then,*

- *(i)With probability at least* 1 − *δ, each s_i_ can be identified through querying* 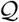 *if* 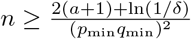
- *(ii) With probability at least* 1 − *δ the L1 distance between the predicted abundances* 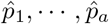 *by setting* 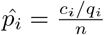 *and the true (unnormalized) abundances* 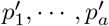 *is at most ϵ if* 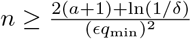.
- *(iii) Given n such reads in a query, with probability at least* 1 − *δ, the L1 distance between the predicted abundances* 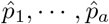 *by setting* 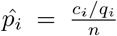 *and the true (unnormalized) abundances* 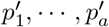 *is bounded by* 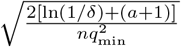. *where c_i_ denotes the number of reads assigned to s_i_*.

See Supplementary Methods for a proof.

## 3 Results

In order to evaluate the overall running time, memory utilization and accuracy of `CAMMiQ`, we have established three datasets to be indexed, all based on NCBI’s RefSeq database [57]. The first dataset aims to evaluate the species level performance of `CAMMiQ` and thus includes one representative genome from each bacterial species. The next dataset aims to evaluate `CAMMiQ`’s strain level performance by including all strain level genome data from RefSeq. Since RefSeq database is biased towards a handful of genera (e.g. *Salmonella* genus is represented by more than 300 strains) we also generated a subspecies level dataset where each species is represented by only a handful of strains as described in detail below.

On the three index datasets, we evaluated `CAMMiQ` on both simulated and real bacterial read sets. The simulated query sets aimed to measure the comparative performance of `CAMMiQ` against some of the best used methods available. The real query sets aimed to evaluate the accuracy of `CAMMiQ` on metatranscriptomic reads from single human immune cells deliberately infected with two distinct strains of the intracellular bacterium *Salmonella enterica* [58].

For each dataset to be indexed and each corresponding query set we compared `CAMMiQ`’s performance with some of the best computational methods available. As will be demonstrated, `CAMMiQ`’s performance is superior to all alternatives in almost all scenarios we tested.

Below, in Section 3.1 we give a detailed description of the index data sets, simulated and real query sets we used, as well as the alternative computational methods we tested to benchmark `CAMMiQ`’s performance. Then in Section 3.2, we demonstrate the advantage offered by `CAMMiQ`’s (i) utilization of doubly-unique substrings – in addition to unique substrings, and (ii) consideration of shortest such substrings instead of fixed length *k*-mers as per CLARK. In Sections 3.3 and 3.4 we demonstrate `CAMMiQ`’s comparative accuracy against alternative metagenomic analysis methods on simulated query sets through the use of our species-level and strain-level datasets. `CAMMiQ`’s comparative performance on computational resources is then demonstrated in Section 3.5. Finally in Section 3.6 we demonstrate `CAMMiQ`’s performance on real metatranscriptomic query sets through its use of our the subspecies-level dataset. The results of `CAMMiQ` in this setup was compared against that of GATK PathSeq which was used for the same purpose earlier [14], as well as BLASTN [9].

### 3.1 The Experimental Setup: Index Datasets, Queries, Benchmarked Methods, Hardware

Our “species-level” index dataset consists of all complete bacterial genomes in NCBI’s RefSeq [57] Database (downloaded on 06/16/2019). We (randomly) selected one representative genome per species out of 13737 reference genomes representing 4122 distinct species. This resulted in a total of *m* = 4122 genomes with a total length of *M* = 3.4 * 10^10^ (including the reverse complement contigs). On this index dataset we used simulated queries only - as will be described below. Since our goal here is to measure `CAMMiQ`’s relative performance against available methods, we also benchmarked state of the art *k*-mer based and marker-gene based metagenomics profiling tools on this index dataset. Specifically we used Kraken2 [59] (the latest version of Kraken [12]), KrakenUniq [44], CLARK [26] and MetaPhlAn2 [60]; these four tools provide very similar functionality to `CAMMiQ` such as read level classification and abundance estimation.

Our “species-level” index dataset consists of all complete bacterial genomes in NCBI’s RefSeq [57] Database (downloaded on 06/16/2019). We (randomly) selected one representative genome per species out of 13737 reference genomes representing 4122 distinct species. This resulted in a total of m = 4122 genomes with a total length of *M* = 3.4 * 10^10^ (including the reverse complement contigs). On this index dataset we used simulated queries only - as will be described below. Since our goal here is to measure `CAMMiQ`’s relative performance against available methods, we also benchmarked state of the art *k*-mer based and marker-gene based metagenomics profiling tools on this index dataset. Specifically we used Kraken2 [59] (the latest version of Kraken [12]), KrakenUniq [44], CLARK [26] and MetaPhlAn2 [60]; these four tools provide very similar functionality to `CAMMiQ` such as read level classification and abundance estimation.

Our “strain-level” index dataset is smaller: it is restricted to 614 Human Gut related genomes (according to [61]) and is designed to evaluate `CAMMiQ`’s strain level identification and quantification performance against the available tools mentioned above. Again, we used simulated query sets for this index dataset. Note that on both the strain-level and species-level index datasets, we evaluated `CAMMiQ` on the most general 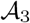 type queries.

Perhaps our most interesting results are on the “subspecies-level” index dataset, which consists of 3395 selected genomes from the 13737 complete bacterial genomes; this dataset was primarily designed to evaluate `CAMMiQ`’s accuracy on query sets involving real metatranscriptomic samples obtained from 262 single human immune cells [58], each exposed to or infected with one of the two distinct *Salmonella* strains. In addition there are 80 uninfected cells used as negative controls. The reads from each cell forms a natural query set for this index dataset. A recent study [14] used the GATK PathSeq tool [10] in order to validate the *Salmonella* strains associated with each cell with limited success. Unlike the tools benchmarked above, GATK PathSeq is alignment based; as a consequence it is slower than the above alternatives but is expected to be more accurate. Interestingly `CAMMiQ`’s ability to distinguish cells exposed to or infected with specific strains of *Salmonella* was better than PathSeq’s ability to do the same - with the added bonus that it is much faster. Note that on metatranscriptomic query sets we used query types 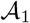 and 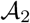 rather than the most general query type 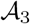.

In order to systematically compare the performance of `CAMMiQ` and other metagenomic profiling tools, we generated several simulated metagenomes as query sets, summarized in Table 1. The upper part of the table corresponds to simulated data from our species-level dataset and the middle part corresponds to our strain-level dataset.

**Table 1:**
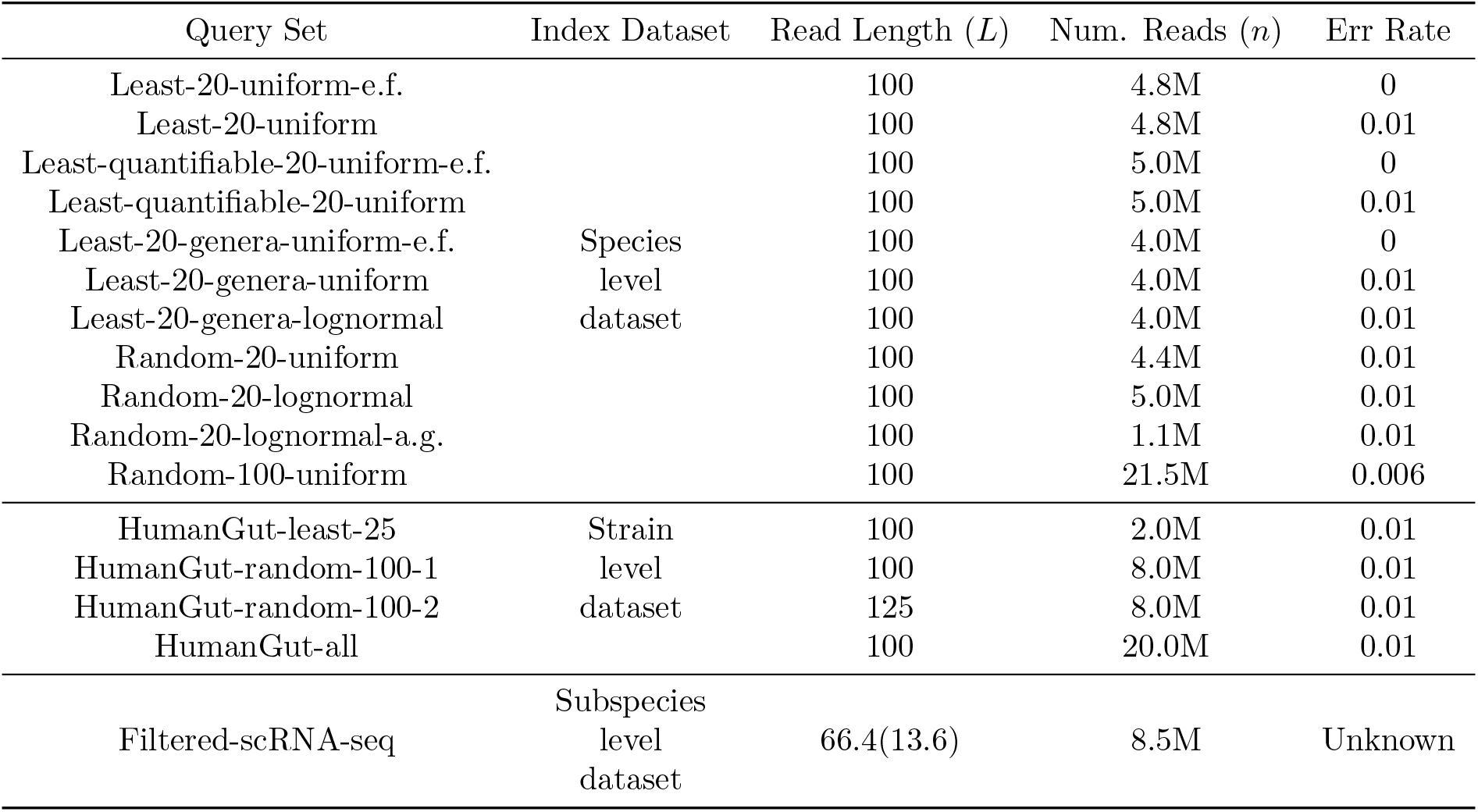
Synthetic and real bacterial read sets used to benchmark `CAMMiQ`’s performance against the best available metagenomic classification and abundance estimation tools. The top collection of queries involve genomes from our species-level index dataset consisting of 4122 distinct bacterial species. The middle collection were sampled from our strain-level index dataset of 617 gastrointestinal associated bacteria (possibly incompletely assembled) provided in [61]. The bottom collection were real single cell RNA-seq reads sequenced from 342 immune cells infected with *Salmonella enterica* [58]; the corresponding subspecies-level index dataset consists a selection of 3395 bacterial genomes from 2753 distinct species.

The first set of simulated metagenomes aim to assess how well `CAMMiQ` identifies species in a query. For that we simulated a metagenome consisting of the 20 genomes that have the lowest number of unique *L*-mers in our species-level dataset. Each genome in the mixture was simulated to have similar read coverage. The very first query we generated from this mixture (denoted Least-20-uniform-e.f.) had no read errors. The second query (denoted Least-20-uniform) had i.i.d. substitution errors occurring at a rate of 1%. Note that the 20 genomes we used in this mixture are intrinsically difficult to be identified by CLARK and other tools we compared. However since these genomes have many doubly-unique *L*-mers - which are sometimes shared with multiple other genomes, they could be identified by `CAMMiQ` (see Supplementary Methods and Figure 5 for a more detailed explanation).

The second set of metagenomes we simulated aim to assess the species-level quantification performance of `CAMMiQ`. This simulated metagenome consisted of the 20 genomes which are among the 50 genomes in our species-level dataset with the lowest proportion of unique *L*-mers, but had the highest proportion of doubly-unique *L*-mers, making them somewhat easier to identify in comparison to the simulated metagenome above, but difficult to quantify by tools other than `CAMMiQ`. We again generated two simulated read collections from this mixture, one with no sequencing errors (denoted Least-quantifiable-20-uniform-e.f.) and another with i.i.d. substitution rate of 1% (denoted Least-quantifiable-20-uniform). The ability of `CAMMiQ`’s ILP formulation to simultaneously determine the presence and abundance of genomes in these queries help it outperform the alternatives. (See Supplementary Methods and Figure 5 for a more detailed explanation.)

Even though RefSeq identifies each genome in our species-level dataset to represent a distinct species, a few of them have “unclassified” lineages at the species level. (Some of the genomes in the above queries are among them; see Figure 5 in the Supplementary Methods.) For example, *Rhizobium sp. N1314* with Taxonomy ID: 1703961 has Rank: species; however its Lineage is noted as *unclassified Rhizobium.* Because of this ambiguity, we generated a third set of metagenomes, again consisting 20 of those 50 genomes with the lowest proportion of unique *L*-mers, this time making sure that each of these 20 genomes represent a distinct genus. We generated 3 queries from this set of genomes. The first one had uniform coverage and had no sequencing errors (denoted Least-20-genera-uniform-e.f.). The second and third both had i.i.d. substitutions at a rate of 1%; the second had a uniform read coverage (denoted Least-20-genera-uniform), while the third had lognormal distribution (denoted Least-20-genera-lognormal).

In addition to the above particularly challenging queries, we simulated a number of additional read collections from 20 to 100 randomly chosen genomes from our species-level dataset. Unlike the above queries, all these read collections had i.i.d. substitution errors; the first three queries at a rate of 1% and the last query at a rate of 0.6%. Among them, the first simulated query (denoted Random-20-uniform) included reads from 20 genomes, each with similar read coverage. The second (denoted Random-20-lognormal) again included reads from 20 genomes, this time with coverages obeying a log-normal distribution. The third (denoted Random-20-lognormal-a.g.) included reads from 20 genomes, again with log-normal coverage distribution; what makes this query unique is that 10% of the reads were from an additional genome (denoted in the dataset name as a.g.) not included in our species-level dataset and thus is not part of `CAMMiQ`’s index. The fourth (denoted Random-100-uniform) included reads from 100 randomly chosen genomes from our species-level dataset, all with similar coverage.

In order to assess `CAMMiQ`’s performance in strain level identification and quantification, we simulated queries involving genomes from a database of 614 strains of human gastrointestinal bacteria [61]^3^ from 409 species. We again simulated multiple queries, the first one involving reads from 25 strains with the smallest number of unique *L*-mers (denoted HumanGut-least-25), next two involving reads from randomly selected 100 strains, the first with *L* = 100 as per the remainder of the queries (denoted HumanGut-random-100-1), and the second with *L* = 125 (denoted HumanGut-random-100-2), and the final involving reads sampled from 409 randomly picked strain level genomes (denoted HumanGut-all), each representing a distinct species in the dataset. Note that none of these queries included more than one strain per species since two distinct strains from a species are not likely to be simultaneously present in a metagenomic sample.

As mentioned earlier, the index we built to respond to these queries consisted of all the 614 strain level genomes described above. The majority of these genomes are not complete and is comprised of multiple contigs; we filtered out any contig with length < 10KB and built the index on the remaining contigs. This resulted in seven strains without a single unique 100-mer and one strain without a single unique or doubly unique 100-mer. This last genome of *Bacillus andreraoultii* was excluded from our queries since it contains no indexable substring.

In our final experiment, we applied CAMMiQ to a “gold-standard” query set consisting of immune cells infected *ex vivo* with an *intracellular* bacteria *Salmonella enterica* and subsequently sequenced using singlecell RNAseq (scRNA-seq) [58] to validate its feasibility of identifying microbial reads from real datasets. Specifically, this query, denoted as Filtered-scRNA-seq and also summarized in Table 1, consists of 342 monocyte-derived dendritic cells (moDCs) infected with either the *D23580* (STM-D23580) or the *LT2* (STM-LT2) strain of *Salmonella enterica* and sequenced using Smart-seq2 platform. The corresponding index we built to respond to this query consisted of the sparsified set of unique and doubly unique substring (with *L* = 75, *L*_min_ = 26 and *L*_max_ = 50) from the subspecies-level dataset of 3395 selected complete bacterial genomes in NCBI’s RefSeq Database. To create this subspecies-level dataset, we first identified 2753 out of the total 4122 species to which at least 10 reads were mapped using GATK PathSeq [10] - and then we subsampled the 4325 (out of the total 13737) genomes from these 2753 species by only keeping one representative for each child of a species level taxonomic ID in NCBI’s Taxonomy Database; The only exception is we replaced the sampled genome of *Salmonella enterica subsp. enterica* (Taxonomy ID: 59201) with the genome of the two related strains *D23580* (Taxonomy ID: 568708) and *LT2* (Taxonomy ID: 99287), which results in the final 3395 subspecies level representatives. Note that as a final step (which is in contrast to the index used in the above simulated queries), we removed all plasmids contained in these genomes, since the plasmids in many genomes downloaded from RefSeq were missing, which may cause potential false positive “unique” or “doubly unique” substrings.

All of our experiments were run on a Linux server equipped with 40 Intel Xeon E7-8891 2.80 GHz processors, with 2.5 TB of physical memory and 30 TB of disk space. The ILP solver used by `CAMMiQ` is IBM ILOG CPLEX 12.9.0.

### 3.2 Comparative Utility of Variable-Length and Doubly-Unique Substrings

In this section we demonstrate the theoretical advantage offered by `CAMMiQ` in comparison to other tools, due to its utilization of not only unique *k*-mers (as per CLARK), but substrings of any length, which could be either unique or doubly-unique - within the species-level dataset we constructed. For that we compared the proportion of *L*-mers from each genome *s*_*i*_ in this dataset (for read length *L* = 100) that are unique or doubly-unique (and thus can be utilized by `CAMMiQ`) with the proportion of *L*-mers that include a unique *k*-mer (that can be utilized by CLARK) for *k* = 30.

Figure 2A summarizes our findings: on the horizontal axis, the genomes are sorted with respect to the proportion of unique and doubly-unique *L*-mers they have; the vertical axis depicts this proportionality (from 0.0 to 1.0). The four plots we have are for the proportion of unique *L*-mers, doubly-unique *L*-mers, the combination of unique and doubly-unique *L*-mers (all utilized by `CAMMiQ`), and the *L*-mers that include a unique *k*-mer (utilized by CLARK). As can be seen, roughly three quarters of all genomes in this dataset (and thus many of the bacterial species) are easily distinguishable since a large fraction of their *L*-mers include a unique *k*-mer. However, roughly a quarter of the genomes in this dataset can benefit from the consideration of doubly-unique substrings, especially when their abundances are low. In particular, 66 of these 4122 genomes/species have extremely low proportion of (≤ 1%) unique 100-mers. In fact, the species *Francisella sp. MA06-7296* does not have a single unique 100-mer and the species *Rhizobium sp. N6212* does not have any 100-mer that include a unique 30-mer (in fact any substring of length ≤ *L*_max_ = 50). These two species cannot be identified by CLARK in any microbial mixture, independent of the abundance values.

**Figure 2:**
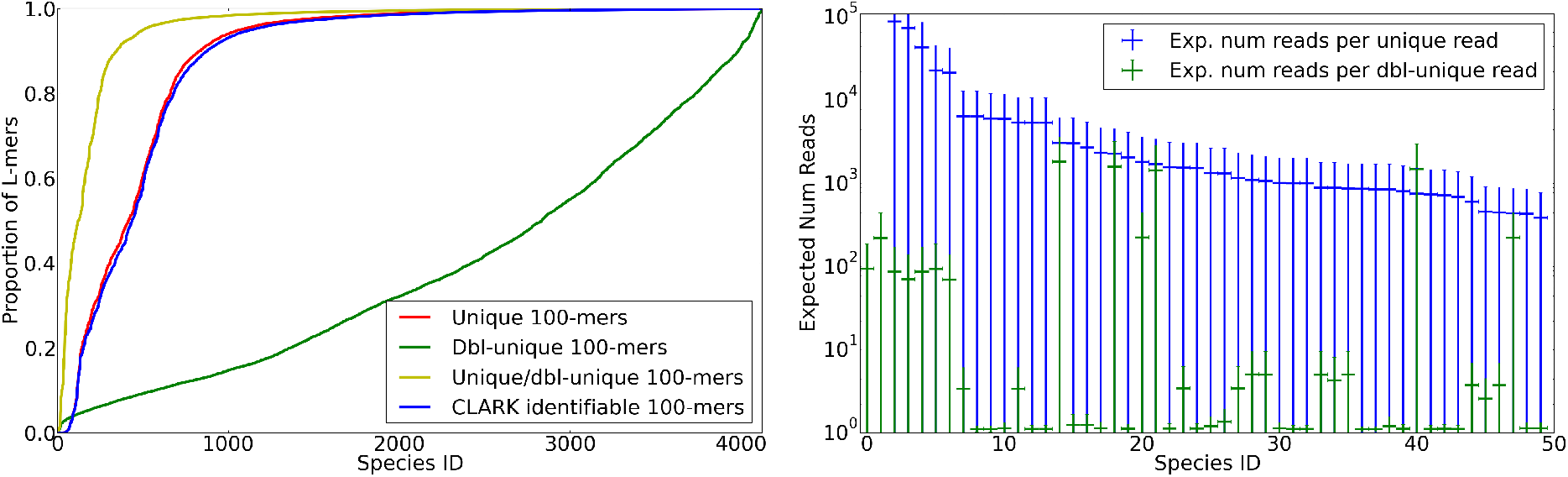
(A) Proportion of *L*-mers (for *L* = 100) that include a unique substring (plotted in red), a doubly-unique substring (plotted in green), or either a unique or a doubly-unique substring (plotted in yellow) in the large dataset of 4122 bacterial genomes from NCBI RefSeq database; these *L*-mers, when presented as reads are utilizable by `CAMMiQ` (when *L*_max_ = 100). Also included are the proportion of *L*-mers that include a unique *k*-mer (*k* = 30) and thus are utilized by CLARK. For each plot, the genomes are independently sorted with respect to the corresponding proportions in ascending order. (B) The expected number of reads (of length *L* = 100) needed to capture one read containing a unique substring (in blue) as well as one read containing a doubly-unique substring (in green) in the 50 genomes with the lowest proportion of unique *L*-mers. Also included are the range for each value within the corresponding standard deviation. Note that for this figure *L*_max_ = 50.

Figure 2B depicts the inverse proportionality of doubly-unique *L*-mers in comparison to unique *L*-mers among 50 genomes that have the lowest proportion of unique *L*-mers - for *L* = 100. The inverse-proportionality of unique or doubly-unique *L*-mers for a genome corresponds to the number of reads to be sampled (on average) from that genome to guarantee that the sample includes one read that is guaranteed to be assigned to the correct genome. In the absence of read errors, this guarantees the correct identification of the corresponding genome in the query. Note that, in half of these 50 genomes, almost all *L*-mers are doubly-unique. This implies that any query involving one or more of these genomes would unlikely to be resolved with CLARK. Yet this query could be handled by `CAMMiQ` since if a read contains a unique or doubly-unique substring, then it can be correctly assigned to the corresponding species as described in Section 2.2.

Note that 3296 of the 4122 species in our species-level dataset have ≥ 90% of their 100-mers as unique. Only a few of these unique substrings do not include a unique 30-mer and thus will be missed by CLARK. This implies that from the accuracy point of view on these genomes, CLARK’s use of *k*-mers instead of the shortest substrings does not put it at a disadvantage on these genomes - when *k* = 30 and *L* = 100. However, as it will become clear later (see Table 5) `CAMMiQ`’s use of shortest substrings, combined with its subsampling strategy gives it an advantage over CLARK (as well as KrakenUniq) with respect to the size of the index structure - despite the fact that `CAMMiQ` needs to index not only unique but also doubly-unique substrings from each genome.

### 3.3 Classification and Quantification Performance at the Species Level

We tested `CAMMiQ` on read collections sampled from the species-level dataset and compared its practical performance with the select metagenomic profiling tools. Perhaps the most widely-used performance measures to benchmark metagenomic classifiers are the proportion of reads correctly assigned to a genome among (i) the set of reads assigned to some genome, i.e. *precision,* and (ii) the full set of reads in the query, i.e. *sensitivity* [26]. In Table 2A we report the precision values for all tools we benchmarked with the exception of MetaPhlAn2, since it does not report individual read assignments. In Table 2B, instead of reporting sensitivity, we report the total number of reads assigned to a genome, since the main goal of Kraken2, KrakenUniq and CLARK is to classify as many reads as possible correctly. The number of reads assigned to a genome (part B of the Table) multiplied by precision (part A of the table) indeed gives this number, i.e. that of correctly classified reads. Note that any read assigned to a taxonomic level higher than (and not including) species level by Kraken2 or KrakenUniq are considered to be not assigned. (In a way, this increases their reported precision but decreases their number of assigned reads.) Also note that `CAMMiQ`’s primary goal is not to classify each read but rather identify and quantify genomes in a query. Nonetheless, we still report its intermediate output as follows. `CAMMiQ` considers certain reads as conflicting; here we consider them as not assigned. It assigns certain reads to a single genome; we consider each such read assigned, and if the assignment is correct, also correctly assigned. `CAMMiQ` then assigns each remaining read ambiguously to two potential genomes.^4^ We consider this read assigned, and in case one of these two genomes is correct, also correctly assigned. Finally since MetaPhlAn2 has an index based on a predetermined database, it is not easy to evaluate this tool’s precision, since the IDs of the genomes indexed by the other four tools do not always correspond to MetaPhlAn2’s genome IDs. As a result we only report its number of assigned reads but not its precision.

**Table 2:**
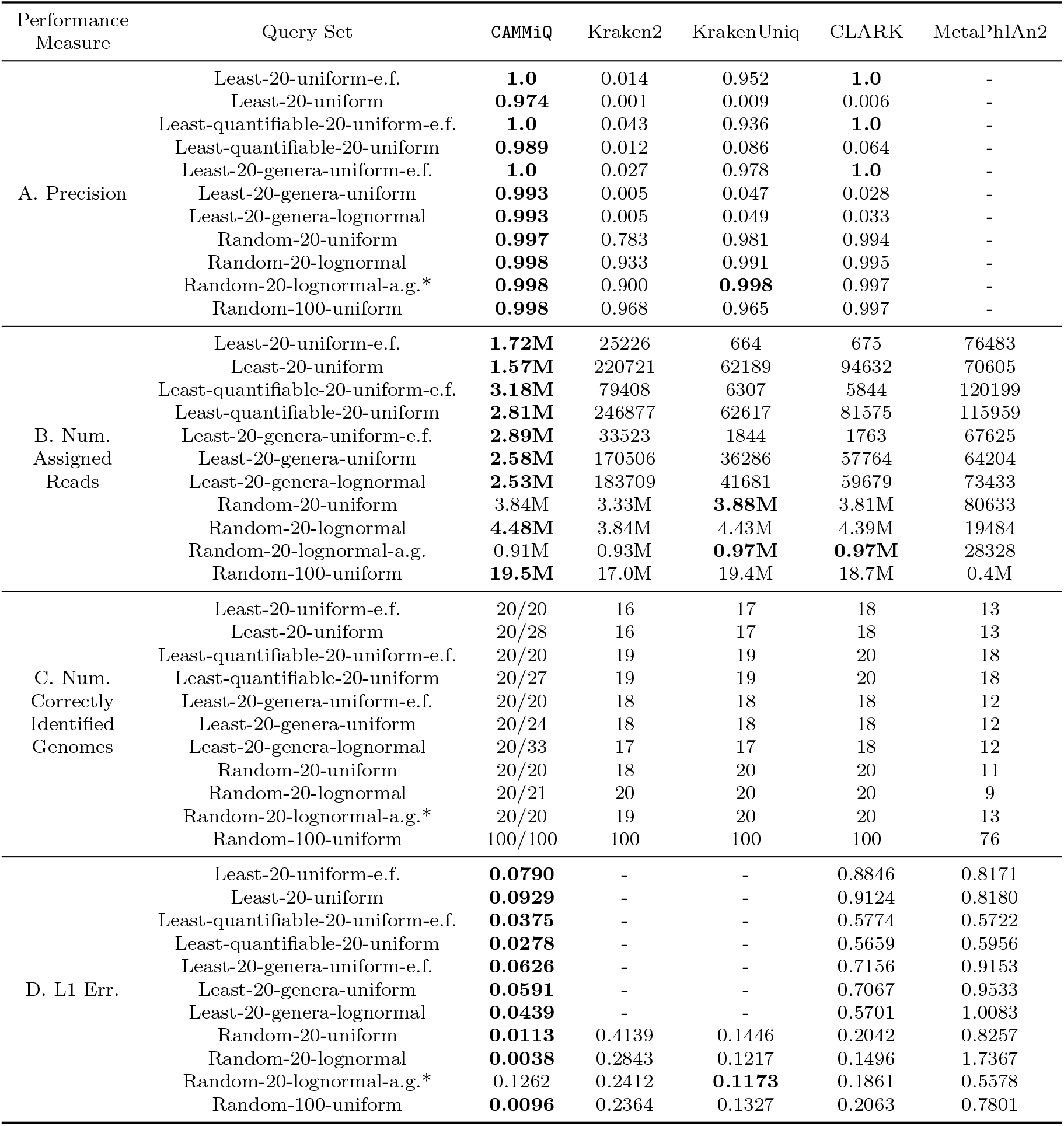
Performance evaluation of `CAMMiQ`, Kraken2, KrakenUniq, CLARK and MetaPhlAn2 on queries from the species level dataset. Precision: the proportion of reads correctly assigned to a genome among the set of reads assigned to some genome (correctly or incorrectly). Number of assigned reads: the total number of reads assigned to some genome. Number of correctly identi_ed genomes: for `CAMMiQ` we report both the number of correctly identi_ed genomes (true positives) and the total number of genomes returned by its ILP formulation (false positive); for Kraken2, KrakenUniq, CLARK and MetaPhlAn2 we only report the number of correctly identified genomes (true positives). Note that we consider MetaPhlAn2 to have a correct identification even if it reports the genus that this genome belongs to. L1 error: the L1 distance between the true relative abundance values (between 0 and 1) and the predicted abundance values for each genome in the query (i.e. positives). We made an exception for MetaPhlAn2, where we measured the genus level L1 distance. Note that we converted the true abundance values reported by Kraken2, KrakenUniq and CLARK by dividing the predicted abundance value for each genome by its length and then normalizing these values by the total abundance value of all genomes. In each of the four measures, the best performing tool's results are highlighted. *: 10% reads in the query Random-20-lognormal-a.g. are from a genome not in the index; any assignment of these reads are necessarily incorrect by all tools except MetaPhlAn2 - which uses its own index, that happens to include this genome.

We benchmarked `CAMMiQ` using its default parameter settings of *L*_min_ = 26 and *L*_max_ = 50, against Kraken2, KrakenUniq and CLARK, with all three using *k*-mer length of 26. All four of these tools used the same genomes for establishing their index. As can be seen in Table 2, `CAMMiQ` demonstrated the best precision for read classification in all of the 11 simulated query sets. With respect to the total number of assigned reads (correctly or incorrectly) on the first 7 queries, i.e. those involving the 20 genomes with the least number of unique *L*-mers (Least-20-uniform-e.f. and Least-20-uniform), those that are the least quantifiable (Least-quantifiable-20-uniform-e.f. and and Least-quantifiable-20-uniform), and those are composed of genomes each from a distinct genus (Least-20-genera-uniform-e.f., Least-20-genera-uniform and Least-20-genera-lognormal), `CAMMiQ` improves over the alternatives not only in terms of precision but also the number of reads assigned, sometimes by an order of magnitude or more. The only exceptions are on those “hypothetical” error-free queries, on which not only `CAMMiQ` and but also CLARK achieves 100% precision. For the above 7 queries Kraken2 has the lowest precision; KrakenUniq has better precision but only to a degree. Furthermore, the number of assigned reads by KrakenUniq is typically the lowest.

On the next four queries, which are easier to identify and quantify, `CAMMiQ`’s peformance is still the best overall. Its precision is the best for all four of these queries while the number of assigned reads are slightly worse than KrakenUniq in two queries. This is likely due to the fact that KrakenUniq produces some incorrect assignments, especially for the Random-20-uniform query on which KrakenUniq has a lower precision. The precision of `CAMMiQ` is identical to KrakenUniq on Random-20-lognormal-a.g. query, where 10% of the reads are sampled from an additional genome not indexed. This is despite that the number of assigned reads from this query is higher for KrakenUniq and CLARK, demonstrating that the introduction of un-indexed species impacts the performance of `CAMMiQ` similarly to the other tools.

We next evaluated the number of correctly identified genomes (for MetaPhlAn2, correctly identified species) in each query, as well as the L1 distance between the true abundance profile and the predicted abundance profile by all five tools on the 11 queries involving our species level dataset. The results can be found in Table 2, parts C and D and Figure 3. Note that `CAMMiQ` and MetaPhlAn2 automatically incorporate a normalization with respect to genome lengths, while Kraken2, KrakenUniq and CLARK simply report the number of reads assigned to each taxonomy rank (for our queries, species) as their abundance profile. In order to compute L1 distances correctly, we converted the true abundance profiles of Kraken2, KrakenUniq and CLARK by dividing the predicted abundance value of each genome by its length and then normalizing these values by the total abundance value of all genomes.

**Figure 3:**
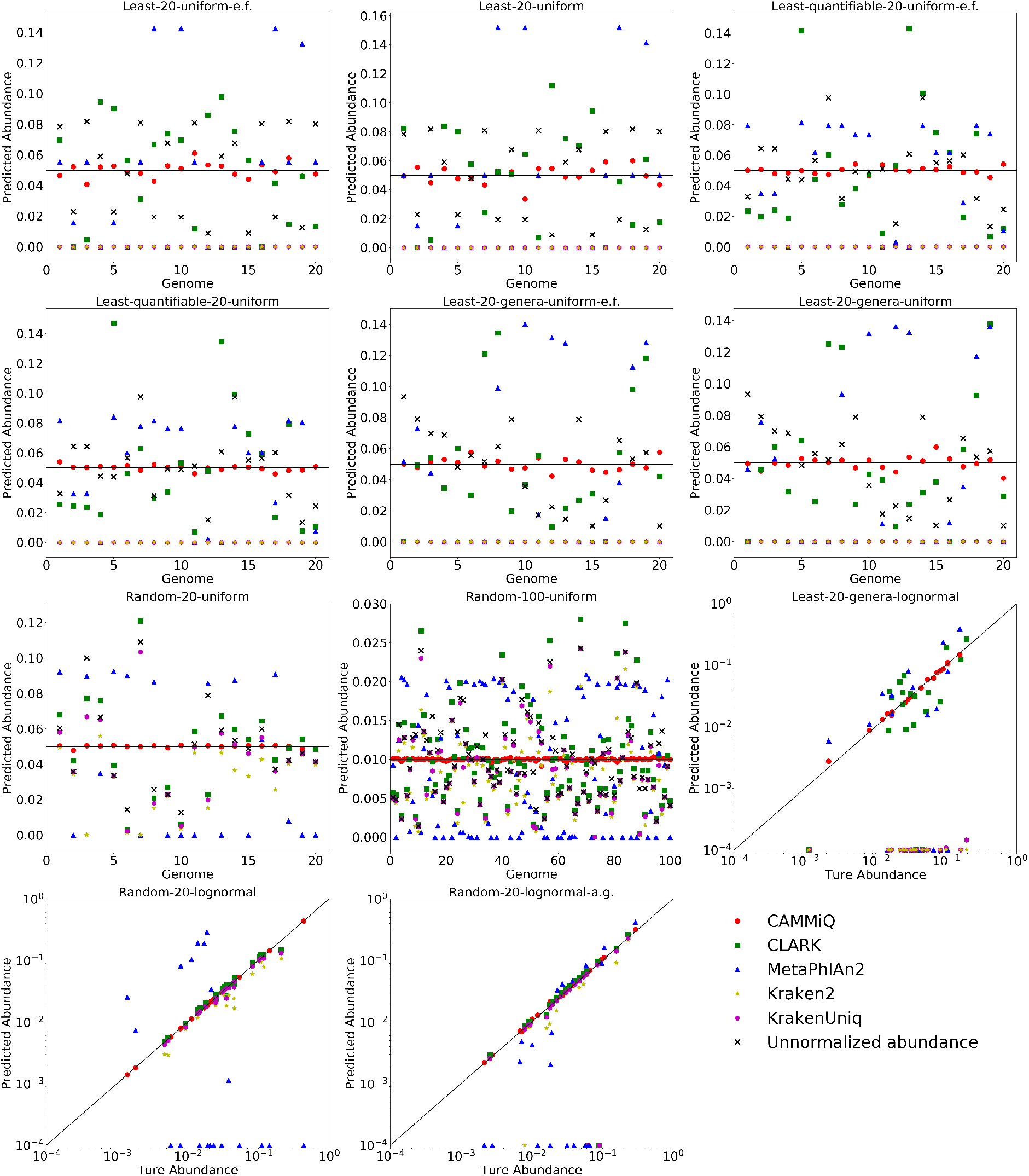
Comparison of `CAMMiQ`’s relative abundance estimates to that of the existing methods for all 11 queries involving the species level dataset. The horizontal axis in the top 8 plots represent the species (i.e., genomes) in an arbitrary order and the vertical axis represent the relative abundance values. The bottom 3 plots are for the queries with log-normal distributions, where the horizontal axis represent the true abundance values while the vertical axis represent the estimated abundance values, both in log-scale.

As can be seen in Table 2 panels C and D, as well as Figure 3, `CAMMiQ` clearly offers the best performance in both identification and quantification. It correctly identified all genomes present in each one of the 11 queries and was not impacted by the additional genome we introduced in the Random-20-lognormal-a.g. query. As importantly, it only returned very few false positive genomes for the most challenging Least-20-uniform, Least-quantifiable-20-uniform and Least-20-genera-uniform as well as Least-20-genera-lognormal queries, and at most one false positive genome for the remaining 7 queries. Other tools had varying levels of false negatives in these 11 queries. Among them CLARK offered the best false negative performance: it only reported false negatives in the queries involving a genome without a single unique *k*-mer. As can be expected, MetaPhlAn2 had the worst performance with respect to false negatives, very likely due to the incompleteness of its marker gene list^5^. This also led to a larger L1 distance than the other tools, even for the relatively easy queries. Kraken2 and KrakenUniq were also prone to have a few false negatives, though less than MetaPhlAn2. Furthermore their predicted abundances are smaller than the true abundance values, because they may assign reads to higher taxonomic ranks than the species level; see Figure 3. This is especially evident in the in the first 7 (difficult) queries: even though Kraken2 and KrakenUniq identified the majority of the genomes correctly, the abundance values they reported on these genomes were all close to 0 and thus the L1 distances turned out to be very close to 1 (see Figure 3). As a result, these values are not reported in Table 2D. Note that we do not report the number of false positive genomes returned by each tool on these 11 queries. This is primarily because of the fact that CLARK, Kraken2 and KrakenUniq are not designed to identify genomes in a query mixture and thus do not “care” whether the false positive assignments are distributed across many genomes (which would result in a large number of false positives) or are concentrated in a few genomes (resulting in a few false positives). MetaPhlAn2 aims to identify genomes however it can do so in any taxonomic level. As such, all these four tools return a very large number of false positives, especially for the first 7 queries, but since we felt that this could be unfair due to the above reasons, we decided to report only the false positives reported by `CAMMiQ`, noting that its genome identification performance is significantly better than the other tools. We ignore the false positive calls also in the L1 measure since it is calculated only on the true positive genomes. This explains the single query and measure for which `CAMMiQ` seems to have performed worse than an alternative, namely KrakenUniq: on Random-20-lognormal-a.g. KrakenUniq’s L1 distance is slightly better than `CAMMiQ`. We remind that in this query, 10% of the reads are sampled from a genome not in the index. Since these reads can not be assigned through any means, the relative abundances of the other genomes will be overestimated by ~ 10%, provided that there are no false positive genomes as per `CAMMiQ`. However, KrakenUniq’s false positive calls (there are several) reduces its relative abundance estimates for the true positive genomes, and gives a seemingly better L1 distance. This can be observed in Figure 3, which depicts for each query, relative abundance estimate for each genome by each one of the tools we benchmarked. As can be clearly seen, `CAMMiQ`’s individual abundance estimates are just on the mark for each one of the genomes even for the most difficult queries.

We finally evaluated the impact of two important parameters for `CAMMiQ`: *α*, the minimum relative read count threshold for reporting a genome, and *L*_min_, the minimum unique or double-unique substring length (values larger than the default value of 50 for *L*_max_ did not have a big impact and thus are not reported here). In Table 3 we report the results for each possible combination of *L*_min_ = 21, 26, 31 and *α* = 0.001,0.0001, on 8 of the 11 queries, omitting the 3 error free queries (on which the impact on precision is minimal). As we increase *L*_min_, `CAMMiQ`’s precision improves, however its read assignment performance deteriorates. Interestingly, its predicted abundance values did not change much with increasing *L*_min_. As a result we set the default *L*_min_ to 26 in `CAMMiQ`. On the other hand, increasing the value of α, decreased the number of false positives in `CAMMiQ`’s output, particularly in the most difficult queries. However, as a result of this, for the queries Least-20-genera-lognormal and Random-20-lognormal-a.g., those genomes with low abundance values were disregarded by `CAMMiQ`, leading to false negatives. `CAMMiQ` allows the user to set the parameter α with prior knowledge on the reads to be queried (e.g., the expected read coverage or the number of genomes in the query); we set its default value to 0.0001.

**Table 3:**
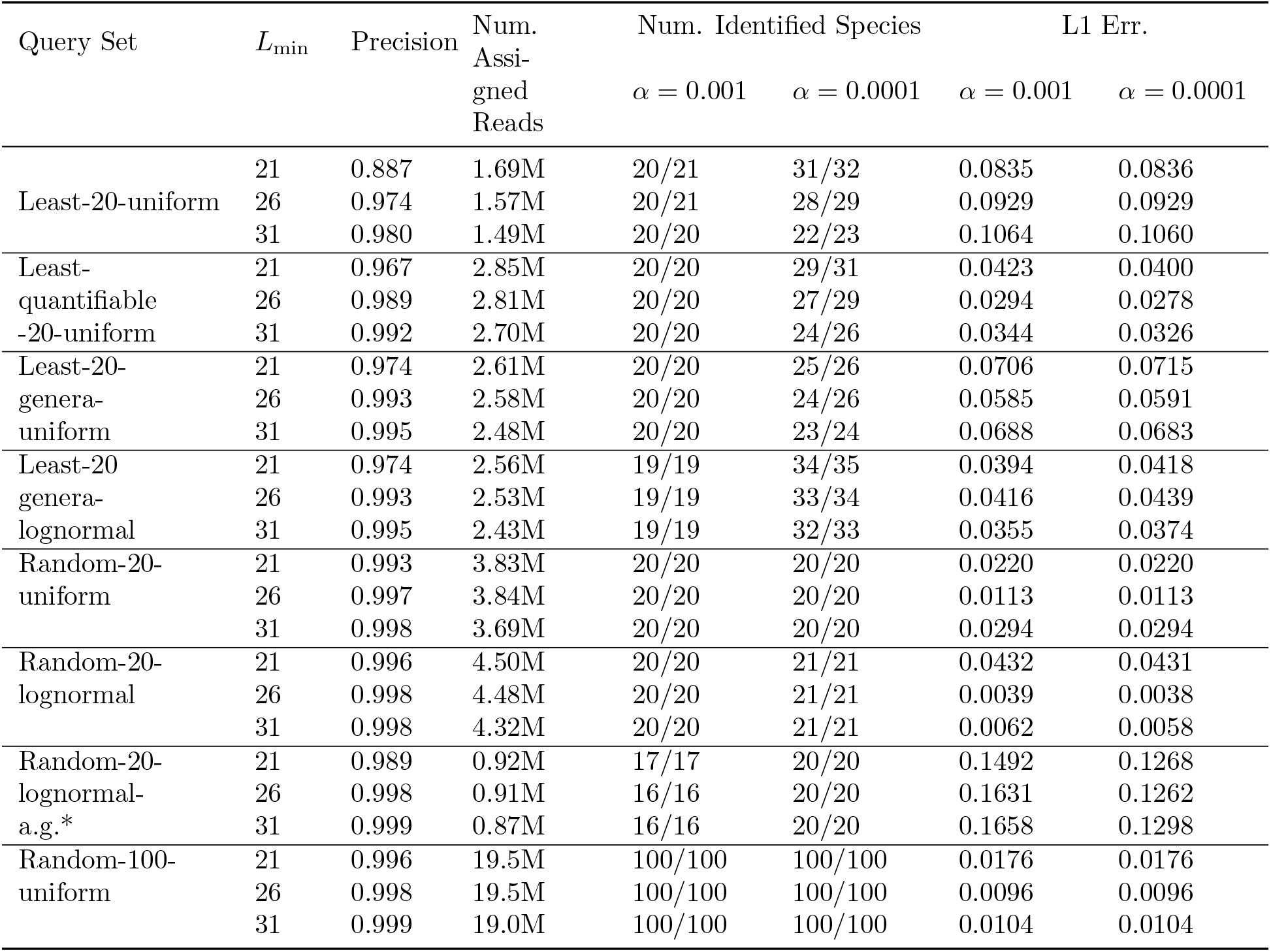
Performance of `CAMMiQ` as a function of minimum unique/doubly-unique substring length *L*_min_ = 21, 26, 31, and minimum relative read count threshold *α* = 0.001, 0.0001 to report a genome. Precision: the proportion of reads correctly assigned to a genome among the set of reads assigned to some genome, correctly or incorrectly. Number of assigned reads: the total number of reads assigned to some genome. Number of identified genomes: the number of genomes returned by `CAMMiQ`’s ILP formulation v.s. the number of genomes that have sufficient read assignments (determined by *α*). L1 error: the L1 distance between the true relative abundance values (between 0 and 1) and the predicted abundance values for each genome in the corresponding query. *: 10% reads in the query Random-20-lognormal-a.g. are from a genome not in the index; any assignment of such a read to a genome is necessarily incorrect.

### 3.4 Quantification Performance at the Strain Level

We finally evaluated `CAMMiQ`’s performance on queries composed from our strain-level dataset that consists of 614 Human Gut related genomes (strains) from 409 species [61] as described in Section 3.1. Three of the tools we evaluated, namely CLARK, Kraken2 and KrakenUniq, simply aim to perform read classification, which we evaluated in Section 3.3. They are not designed to identify or quantify genomes, especially at the strain level. On the other hand MetaPhlAn2’s index have strain level information and through that it attempts to identify and quantify strains. As a result we report on the performances of both `CAMMiQ` and MetaPhlAn2 in Table 4. As can be seen, `CAMMiQ` managed to identify and accurately quantify all strains in the queries HumanGut-random-100-1 and HumanGut-random-100-2, and > 98% strains in the query HumanGut-all, with almost no false positives. Here we only report the strain level calls made by MetaPhlAn2 (it made additional calls at the species level or higher), based on our best attempt to match its calls to the strain IDs available to us. It may be possible to slightly increase the true positive values (the first value) reported here.

**Table 4:**
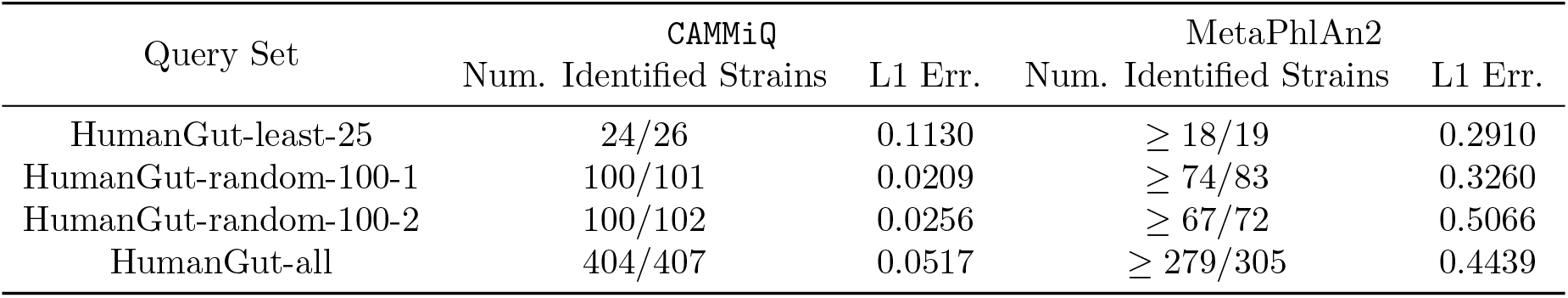
`CAMMiQ`’s quantification performance at strain level, compared to MetaPhlAn2. Number of identified strains: the number of true positive strains/the total number of strains identified. L1 error: the L1 distance between the true relative abundance values and the predicted abundance values, across all strains in the query.

| Query Set | CAMMiQ |  | MetaPhlAn2 |  |
| --- | --- | --- | --- | --- |
|  | Num. Identified Strains | L1 Err. | Num. Identified Strains | L1 Err. |
| HumanGut-least-25 | 24/26 | 0.1130 | $\geq 18/19$ | 0.2910 |
| HumanGut-random-100-1 | 100/101 | 0.0209 | $\geq 74/83$ | 0.3260 |
| HumanGut-random-100-2 | 100/102 | 0.0256 | $\geq 67/72$ | 0.5066 |
| HumanGut-all | 404/407 | 0.0517 | $\geq 279/305$ | 0.4439 |

### 3.5 Performance on Computational Resources

We compared the running time and memory use of `CAMMiQ`, Kraken2, KrakenUniq, CLARK and MetaPhlAn2 in responding to the queries; see Table 5. We do not report the time for building the index or loading it into memory, since this is performed only once - all tools roughly need a couple hours to construct the index on the species level dataset. When it comes to querying time, `CAMMiQ` is outperformed only by Kraken2, typically by a factor of 3. This is primarily due to the fact that Kraken2’s index is much smaller than that of `CAMMiQ` since it consists of a small subset of unique *k*-mers. This compact index structure substantially impacts its performance, which is improved by KrakenUniq, through its consideration of all unique *k*-mers. The resulting index size of KrakenUniq is comparable to that of `CAMMiQ` (`CAMMiQ`’s is slightly higher as it also includes doubly-unique substrings), however its running time is almost twice as much, even though it does not solve an ILP. `CAMMiQ` owes its superior identification and quantification performance to its ILP formulation, however it is not time-wise penalized by it. The best memory footprint is achieved by MetaPhlAn2 through the use of its own index, however both its run time and its identification/quantification performance is worse than the others.

**Table 5:**
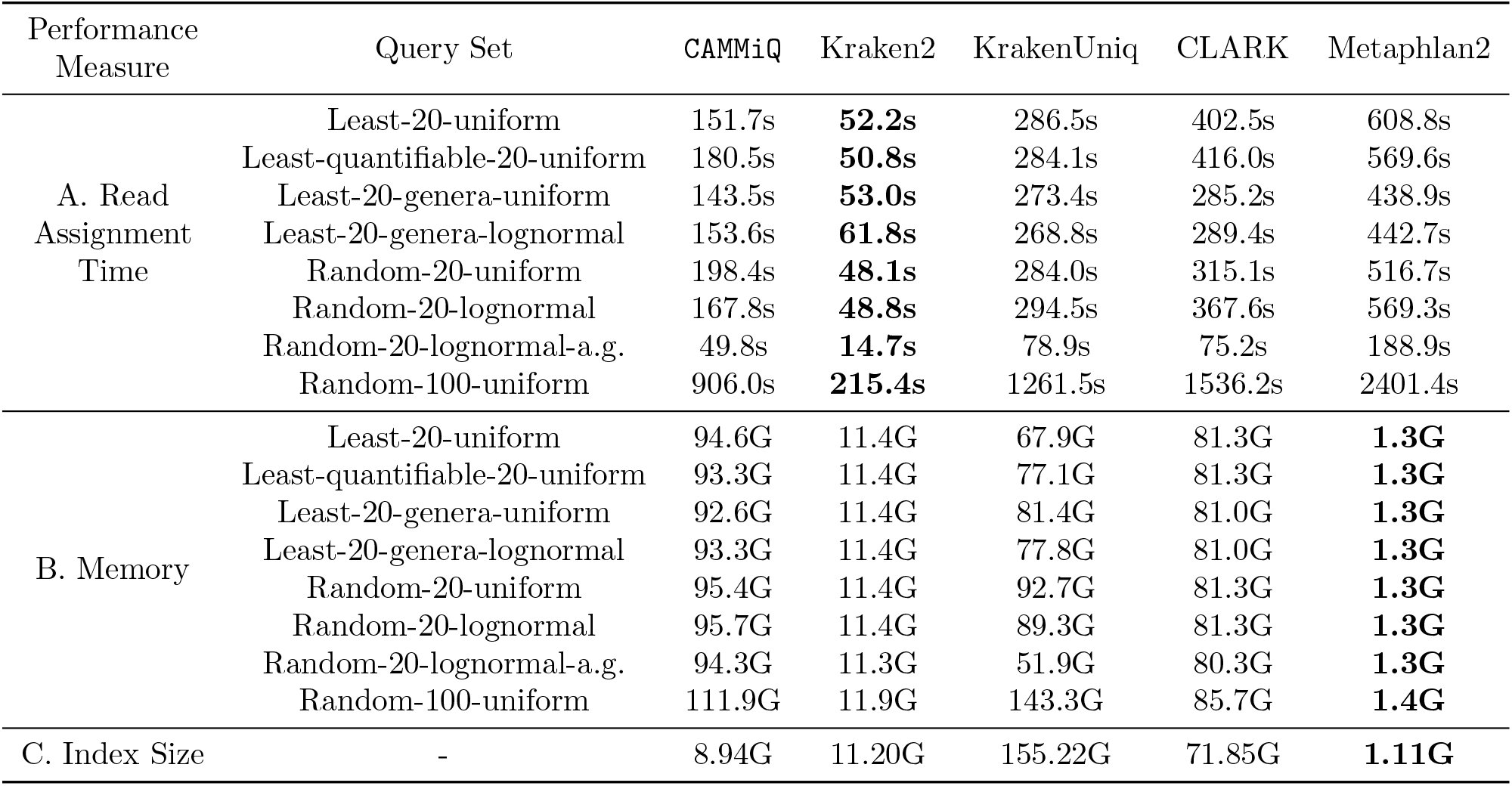
Comparison of the computational resources required by `CAMMiQ`, Kraken2, KrakenUniq, CLARK and MetaPhlAn2, when run on a single thread.

| Performance Measure | Query Set | CAMMiQ | Kraken2 | KrakenUniq | CLARK | Metaphlan2 |
| --- | --- | --- | --- | --- | --- | --- |
| A. Read Assignment Time | Least-20-uniform | 151.7s | **52.2s** | 286.5s | 402.5s | 608.8s |
|  | Least-quantifiable-20-uniform | 180.5s | **50.8s** | 284.1s | 416.0s | 569.6s |
|  | Least-20-genera-uniform | 143.5s | **53.0s** | 273.4s | 285.2s | 438.9s |
|  | Least-20-genera-lognormal | 153.6s | **61.8s** | 268.8s | 289.4s | 442.7s |
|  | Random-20-uniform | 198.4s | **48.1s** | 284.0s | 315.1s | 516.7s |
|  | Random-20-lognormal | 167.8s | **48.8s** | 294.5s | 367.6s | 569.3s |
|  | Random-20-lognormal-a.g. | 49.8s | **14.7s** | 78.9s | 75.2s | 188.9s |
|  | Random-100-uniform | 906.0s | **215.4s** | 1261.5s | 1536.2s | 2401.4s |
| B. Memory | Least-20-uniform | 94.6G | 11.4G | 67.9G | 81.3G | **1.3G** |
|  | Least-quantifiable-20-uniform | 93.3G | 11.4G | 77.1G | 81.3G | **1.3G** |
|  | Least-20-genera-uniform | 92.6G | 11.4G | 81.4G | 81.0G | **1.3G** |
|  | Least-20-genera-lognormal | 93.3G | 11.4G | 77.8G | 81.0G | **1.3G** |
|  | Random-20-uniform | 95.4G | 11.4G | 92.7G | 81.3G | **1.3G** |
|  | Random-20-lognormal | 95.7G | 11.4G | 89.3G | 81.3G | **1.3G** |
|  | Random-20-lognormal-a.g. | 94.3G | 11.3G | 51.9G | 80.3G | **1.3G** |
|  | Random-100-uniform | 111.9G | 11.9G | 143.3G | 85.7G | **1.4G** |
| C. Index Size | - | 8.94G | 11.20G | 155.22G | 71.85G | **1.11G** |

### 3.6 Identification Performance - Subspecies Level Dataset with Real scRNAseq Queries

As mentioned earlier, for the “real” metatranscriptomic dataset, we employed query types 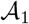 and 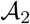 instead of query type 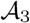. We remind the reader that 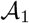 simply requires the computation of all genomes in the index dataset for which there is at least one unique substring observed in the query read set, and 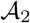 requires the computation of the smallest set of genomes in the index dataset to include all unique or doubly-unique substrings observed in the query read set. Here each query corresponds to the Filtered set of non-human scRNA-seq reads from the corresponding immune cells as described in [58]. More specifically we pre-filtered all query scRNA-seq reads which (i) possibly originate from the human genome, (ii) have low sequence quality and “complexity”, or (iii) map to 16S or 23S ribosomal RNAs on the two *Salmonella* genomes (to avoid incorrect assignment of reads due to “barcode hopping”). When analyzing the resulting genomes, we used annotations provided by [58] and categorized the cells into 5 groups: infected cells that were confirmed to contain intracellular Salmonella (either the STM-LT2 or STM-D23580 strain); bystander cells that were exposed to either the STM-LT2 or STM-D23580 strains but confirmed to not contain intracellular Salmonella; and cells that were mock-infected and sequenced as controls. For each query, we compared the number of reads `CAMMiQ` assigned uniquely to STM-LT2 or STM-D23580 genomes to those aligned (and assigned) by GATK PathSeq [10] as well as BLASTN [9].

As expected, CAMMiQ reconfirmed that the abundance (measured by unique read counts) of *Salmonella* was substantially higher in the infected cells compared to the mock-infected controls; and importantly, the unique STM-LT2 (and STM-D23580) reads were differentially abundant between the cells exposed to or infected with STM-LT2 strain (and STM-D23580 strain respectively; see Figure 4A). Interestingly, CAMMiQ identified a small number of unique STM-LT2 reads in the cells exposed to or infected with STM-D23580 strain and vice versa; these were verified with BLASTN, indicating possible sequencing errors or incorrect assignment of specific reads to cells. Compared with the alignment based approach GATK PathSeq with the same index dataset, CAMMiQ was shown to be more sensitive: on average, it identified (roughly) an order of magnitude higher number of unique STM-LT2 or STM-D23580 reads per corresponding cell. This demonstrates `CAMMiQ`’s potential ability to better identify intracellular organisms at subspecies or strain level; see Figure 4B. ^6^ When it comes to running time, `CAMMiQ` offers several orders of magnitude comparative

**Figure 4:**
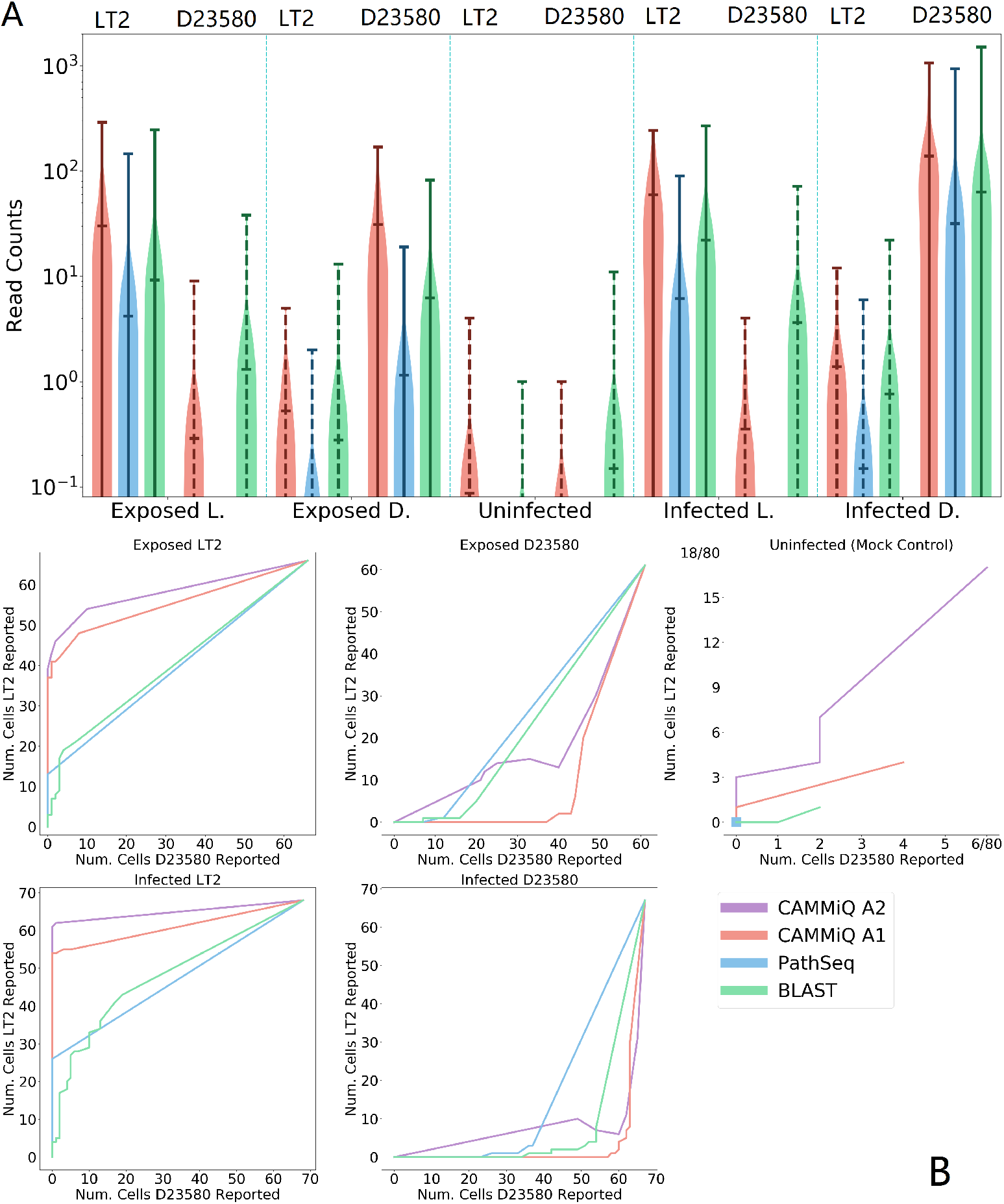
A: Comparison of the distributions of filtered-scRNAseq reads uniquely assigned to the STM-LT2 (the left side in each panel) or STM-D23580 strain (the right side in each panel) by CAMMiQ (red), GATK PathSeq (blue) and BLASTN (green) between the 5 groups of cells that were (i) exposed to STM-LT2; (ii) exposed to STM-D23580; (iii) mock-infected as controls; (iv) infected with STM-LT2; and (v) infected with STM-D23580. Solid lines indicate the strain which the cells have been exposed to or infected with. Ideally the strain with which the cells have been exposed to or infected with should have higher read count values. B: The number of cells with more than *t* reads uniquely assigned to the STM-LT2 strain (y-axis in each panel) or STM-D23580 strain (x-axis in each panel) by CAMMiQ with query type 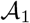 (red), GATK PathSeq (blue) and BLASTN (green), for varying values of *t*. We also included the results of `CAMMiQ` with query type 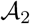 (purple) which uses doubly-unique substrings in addition to unique substrings and thus is not represented in part (A) of the figure. When the threshold *t* is very high, neither strain could be detected by any method in any of the cells - this corresponds to position (0, 0) in each plot. When the threshold is very low (as low as 0), then both strains can be detected in all cells by all tools - this corresponds to position (*c, c*) where *c* corresponds to the total number of cells in a given plot. The plot for an ideal tool should connect positions (0, 0) with (*c, c*) following the axis that represents the particular strain with which the cells are exposed to or infected with, as closely as possible (the mock control is the exception). The plots for `CAMMiQ` are closer to this ideal than PathSeq or BLASTN.

As expected, `CAMMiQ` reconfirmed that the abundance (measured by unique read counts) of *Salmonella* was substantially higher in the infected cells compared to the mock-infected controls; and importantly, the unique STM-LT2 (and STM-D23580) reads were differentially abundant between the cells exposed to or infected with STM-LT2 strain (and STM-D23580 strain respectively; see Figure 4A). Interestingly, `CAMMiQ` identified a small number of unique STM-LT2 reads in the cells exposed to or infected with STM-D23580 strain and vice versa; these were verified with BLASTN, indicating possible sequencing errors or incorrect assignment of specific reads to cells. Compared with the alignment based approach GATK PathSeq with the same index dataset, `CAMMiQ` was shown to be more sensitive: on average, it identified (roughly) an order of magnitude higher number of unique STM-LT2 or STM-D23580 reads per corresponding cell. This demonstrates `CAMMiQ`’s potential ability to better identify intracellular organisms at subspecies or strain level; see Figure 4B. ^6^ When it comes to running time, `CAMMiQ` offers several orders of magnitude comparative advantage on the filtered-scRNAseq queries. Specifically, `CAMMiQ` only took a total of 65.3s for computing 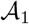 type queries and an additional 2.5s for computing 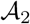 type queries on the entire query set, outperforming GATK PathSeq, which required 29628.1s.

## 4 Discussion

We have introduced `CAMMiQ`, a new computational approach to solve a computational problem that has not been exactly addressed by any available method: given a set 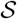 of distinct genomic sequences (of any taxonomic rank), build a data structure so as to identify and quantify genomes in a any query, composed of a mixture of reads from a subset of genomes from 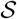. `CAMMiQ` is particularly designed to handle genomes that lack unique features; for that, it reduces the identification and quantification problems to a combinatorial optimization problem that assigns substrings with limited ambiguity (i.e. doubly unique substrings) to genomes so that (in its most general 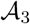 type query), each genome is “uniformly covered”. Because each such substring has limited ambiguity, the resulting combinatorial optimization problem can be very efficiently solved through existing integer program solvers such as IBM CPLEX (and thus are branch-and-bound based, whose run-time performance is determined by the “fan-out” at each decision point). Provided that the doubly-unique substrings of a given genome are not all shared with one other genome, the use of doubly-unique substrings increases `CAMMiQ`’s ability to identify and quantify this genome within a query. As mentioned earlier, in case the dataset to be indexed involves several genomes with high level of similarity, `CAMMiQ`’s data structure and its combinatorial optimization formulation can easily be generalized to include triply or quadruply unique substrings, without much computational overhead.

## 5 Supplementary Methods

### 5.1 Unique substrings from LCP_*u*_ or LCP_*d*_

The pseudocode for the algorithm to compute arrays `SU` and `SD` according to equations (3) and (4) respectively, given `SA`, `LCP`_*u*_ and `LCP`_*d*_ is given below. As can be seen, the algorithm runs in *O(M)* time.

**Algorithm 1.**
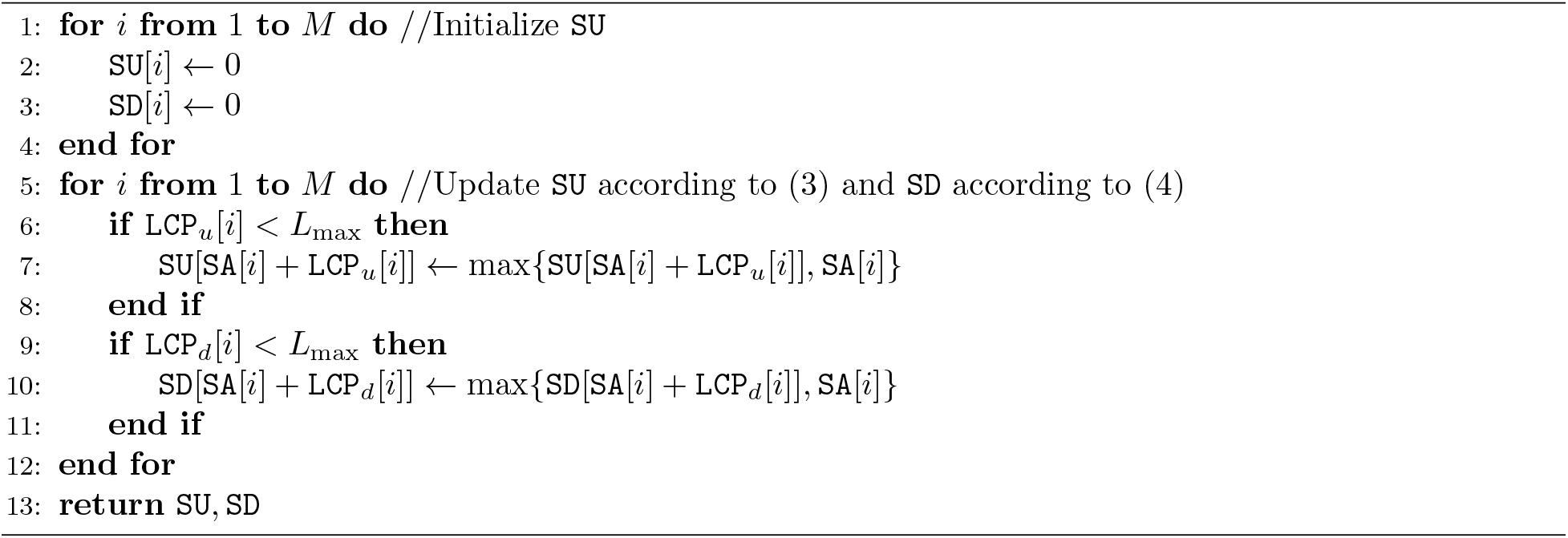
ShortestUniqueFromLCP(`SA`, `LCP`_*u*_, `LCP`_*d*_, *L*_max_)

### 5.2 Computing LCP_*u*_ and LCP_*d*_

In this section we show that `SU` and `SD` can be correctly constructed in *O(M*) time. We start by showing that the definition of `LCP`_*u*_ and `LCP`_*d*_ in Equation (1) and Equation (2) can respectively lead to the shortest substrings occurring in at most one genome or two genomes. Then we give `CAMMiQ`’s detailed implementation of Equation (5) and Equation (6) to compute the `LCP`_*u*_ and `LCP`_*d*_ arrays. Finally we give a running time analysis of this implementation.

First consider the content of `SU` at the end of procedure ShortestUniqueFromLCP. To see the substring *s*[*l : r*] corresponds to the *r*-th entry `SU`[*r*] = *l* (where *l* ≠ 0) in `SU` is unique, meaning it only occurs in genome with ID `GSA`[`SA`^−1^[*l*]], assume that there is another genome *s*_*i′*_ having the same substring *s*[*l′*, *r′*] = *s*[*l : r*] - this leads to a contradiction, since it implies that lcp(`suf`[*l*], `suf`[*l′*]) ≥ *r* − *l* + 1. However, due to the update rule of `SU` and the definition of `LCP`_*u*_, lcp(`suf`[*l*], `suf`[*l′*]) ≤ *r* − *l* for any 1 ≤ *l′* ≤ *M* satisfying `suf`[*l*] and `suf`[*l′*] start on different genomes, namely `GSA`[`SA`^−1^[*l*]] ≠ `GSA`[`SA`^−1^[*l′*]], which is a contradiction. Now, to see *s*[*l : r*] is a shortest unique substring, i.e. no substring of *s*[*l : r*] is unique to genome `GSA`[`SA`^−1^ [*l*]], we show that any *s*[*l : r′* < *r*] and *s*[*l′* > *l,r*] occurs in one other genome *s*_*i′*_. The former case is due to the definition of `LCP`_*u*_ - there exists `suf`[*l′*] on genome *i′* ≠ `GSA`[`SA`^−1^[*l*]] such that lcp(`suf`[*l*], `suf`[*l′*]) ≥ *r* − *l*, implying a substring *s*[*l′* : *l′* + (*r* − *l*) − 1] identical to *s*[*l,r − 1*]; the later case is due to the update rule of `SU` - if *s*[*l*′ > *l,r*] is also unique to genome `GSA`[`SA`^−1^[*l*]], then `SU`[*r*] must be set to *l′* instead of *l*. Therefore, *s*[*l : r*] is a shortest unique substring (to genome with ID `GSA`[`SA`^−1^[*l*]]); on the other hand, if *s*[*l : r*] is a shortest unique substring, then `SU`[*r*] will maintain *l* after `SU` is completely updated.

Now consider the content of `SD` at the end of procedure ShortestUniqueFromLCP. We follow the above proof to show *s*[*l : r*] is a shortest doubly-unique substring (to genome ID `GSA`[`SA`^−1^[*l*]] and possibly another genome *i′*). To see the substring *s*[*l : r*] corresponds to the *r*-th entry `SD`[*r*] = *l* (where *l* ≠ 0) in `SD` occurs in at most two genomes, with ID `GSA`[`SA`^−1^[*l*]] (and possibly *i′*, any genome that `suf`[*l′*] belongs to, giving the largest lcp(`suf`[*l*], `suf`[*l′*])), we can assume there exists a thrid genome *s*_*i′′*_ having the same substring *s*[*l′′*,*r′′*] = *s*[*l,r*] = *s*[*l′,r′*] and similarly obtain a contradiction. Note that according to (2), it is possible to have `LCP`_*d*_[`SA`^−1^[*l*]] ≥ `LCP`_*u*_[`SA`^−1^[*l*]] and in this case *s*[*l : r*] is a unique substring which only occurs in genome `GSA`[`SA`^−1^ [*l*]]. If `LCP`_*d*_[`SA`^−1^[*l*]] < `LCP`_*u*_[`SA`^−1^[*l*]] on the other hand, then *s*[*l : r*] must occur in exactly two genomes, since `SD` is updated according to `LCP`_*d*_ and we can find another suffix of *s* whose length-(*r* − *l* + *1*) (*r* − *l* + 1 ≤ `LCP`_*d*_ [`SA`^−1^[*l*]]) prefix is identical to *s*[*l : r*]. In addition, to see *s*[*l : r*] is a shortest doubly-unique substring, meaning no substring of *s*[*l : r*] occurs only in genome `GSA`[`SA`^−1^[*l*]] and *i′*, we can similarly show that any *s*[*l : r′* < *r*] and *s*[*l′* > *l, r*] can be found in a third genome *s*_*i′′*_, regardless whether `LCP`_*d*_[`SA`^−1^[*l*]] = `LCP`_*u*_[`SA`^−1^[*l*]] or `LCP`_*d*_[`SA`^−1^[*l*]] < `LCP`_*u*_[`SA`^−1^[*l*]].

As a result of the above observations we can now formally state the following lemma.

#### Lemma 2.

*(i)After updating `SU` according to (3), `SU`*[*r*] = *l* ≠ 0 *implies that s*[*l : r*] *is a shortest unique substring to genome `GSA`[`SA`*^−1^[*l*]];

*(ii)After updating `SD` according to (4) `SD`*[*r*] = *l* ≠ 0 *implies that s*[*l : r*] *is a shortest doubly-unique substring to genome `GSA`[`SA`*^−1^ [*l*]] *and i′ = `GSA`[`SA`*^−1^[*l′*]] *where `suf`*[*l′*] *gives the largest lcp(`suf`*[*l*], *`suf`*[*l′*])).

Furthermore, it’s also clear that all unique substrings *s*[*l : r*] are stored in `SU` and all doubly-unique substrings *s*[*l : r*] are stored in SD, as we have considered the suffix of s starting with every possible *l*.

Both `LCP`_*u*_ and `LCP`_*d*_ can be modified to incorporate the minimum length constraint *L*_min_ on unique/doubly-unique substrings. By setting `LCP`_*u*_[*i*] to max{*L*_min_ − 1, `LCP`_*u*_[*i*]} for each entry 1 ≤ *i* ≤ *M*, the corresponding substrings maintained in `SU` should also be unique, and with minimum length *L*_min_. One should be careful however when dealing with `LCP`_*d*_: if `LCP`_*u*_[*i*] ≤ *L*_min_ − 1, then the corresponding substring *s*[`SA`[*i*], `SA`[*i*] + `LCP`_*u*_[*i*]] occurs in only one genome. Therefore we set `LCP`_*d*_[*i*] to ∞ (meaning it’s not considered) if `LCP`_*u*_[*i*] ≤ *L*_min_ − 1 or `LCP`_*d*_[*i*] ≥ `LCP`_*u*_[*i*]; and to max{*L*_min_ − 1, `LCP`_*d*_[*i*]} otherwise (this can be done by first set each `LCP`_*u*_[*i*] to max{*L*_min_ − 1,`LCP`_*u*_[*i*]} and `LCP`_*d*_[*i*] to max{*L*_min_ − 1,`LCP`_*d*_[*i*]}, and then set each `LCP`_*d*_[*i*] to ∞ if `LCP`_*d*_[*i*] ≥ `LCP`_*u*_[*i*]), which ensures the corresponding substrings maintained in `SD` are doubly-unique, and with minimum length *L*_min_.

With the correctness of (3) and (4) in mind, our next concern is how to actually compute `LCP`_*u*_ and `LCP`_*d*_ based on their definitions. In the following we show that (5) and (6) correctly implement (1) and (2), without considering the borderline cases (i.e., for *i* = 1 or *i* = *M*; to handle these cases we can set `GSA`0. = `GSA`[*M* + 1] = 0 and ignore *i*^2−^ when *i* = 1 and *i*^2+^ when *i* = *M*).

#### Lemma 3.

*For any 1 ≤ i ≤ M*,

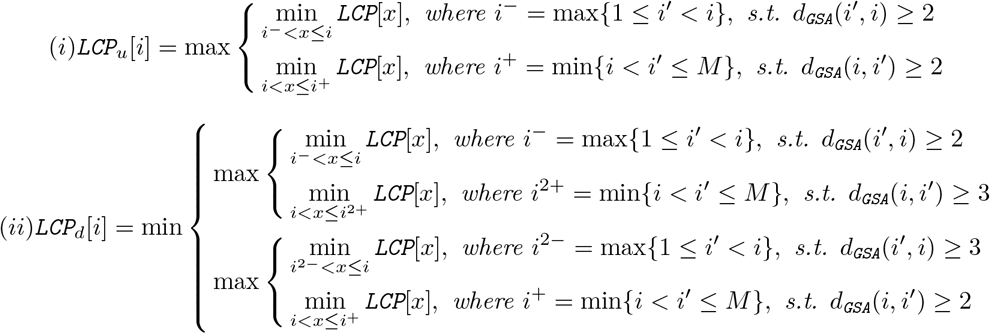

*where d*_GSA_(*i*_1_, *i*_2_) = |{*`GSA`*[*i*_1_], ···, *`GSA`*[*i*_2_]}|.

*Proof.* Let lcp(*i,j*) denote lcp(`suf`[*i*], `suf`[*j*]) for short. We utilize the properties of `SA` and `LCP` array: (a) the longest common prefix of two suffices `suf`[*i*] and `suf`[*j*] (assume `suf`[*i*] is lexicographically smaller than `suf`[*j*]) is lcp(*i, j*) = min{`LCP`[*x*] | *x* ∈ [`SA`^−1^[*i*] + 1, `SA`^−1^[*j*]]}; also we have (b) lcp(*i, j*) ≥ lcp(`SA`[`SA`^−1^ [*i*] − 1], *j*) and lcp(*i, j*) ≥ lcp(*i*, `SA`[`SA`^−1^[*j*] + 1]). (i) follows immediately from these properties:

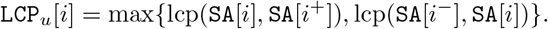

To see (ii), we consider three cases:

- If lcp(`SA`[*i*], `SA`[*i*^+^]) = lcp(`SA`[*i*^−^], `SA`[*i*]), then `LCP`_*d*_[*i*] = lcp(`SA`[*i*], `SA`[*i*^+^]) = lcp(`SA`[*i*^−^], `SA`[*i*]), due to (b). (ii) holds in this case by applying (a).
- If lcp(`SA`[*i*], `SA`[*i*^+^]) < lcp(`SA`[*i*^−^], `SA`[*i*]), then LCP_*d*_[i] = max{lcp(`SA`[*i*], `SA`[*i*^+^]), lcp(SA[i^2−^], `SA`[*i*])} due to (b). Also `LCP`_*d*_[*i*] ≤ lcp(`SA`[*i*^−^], `SA`[*i*]) = max{lcp(`SA`[*i*], `SA`[*i*^2+^]), lcp(`SA`[*i*^−^], `SA`[*i*])} since lcp(`SA`[*i*], `SA`[*i*^2+^]) ≤ lcp(`SA`[*i*], `SA`[*i*^+^]) < lcp(`SA`[*i*^−^], `SA`[*i*]). (ii) therefore holds by applying (a).
- If lcp(`SA`[*i*], `SA`[*i*^+^]) > lcp(`SA`[*i*^−^], `SA`[*i*]), then we have similarly `LCP`_*d*_[*i*] = max{lcp(`SA`[*i*], `SA`[*i*^2+^]), lcp(`SA`[*i*^−^], `SA`[*i*])} due to (b) and `LCP`_*d*_[*i*] ≤ lcp(`SA`[*i*], `SA`[*i*^+^]) = max{lcp(`SA`[*i*], `SA`[*i*^+^]), lcp(SA[*i*^2−^], `SA`[*i*])} since lcp(`SA`[*i*^2−^], `SA`[*i*]) ≤ lcp(`SA`[*i*^−^], `SA`[*i*]) < lcp(`SA`[*i*], `SA`[*i*^+^]). Again we can see (ii) by applying (a), which complete the proof.

Next we give the pseudocode for the algorithm to update `LCP`_*u*_ and `LCP`_*d*_ based on (5) and (6) respectively, and show that they actually run in linear time.

**Algorithm 2.**
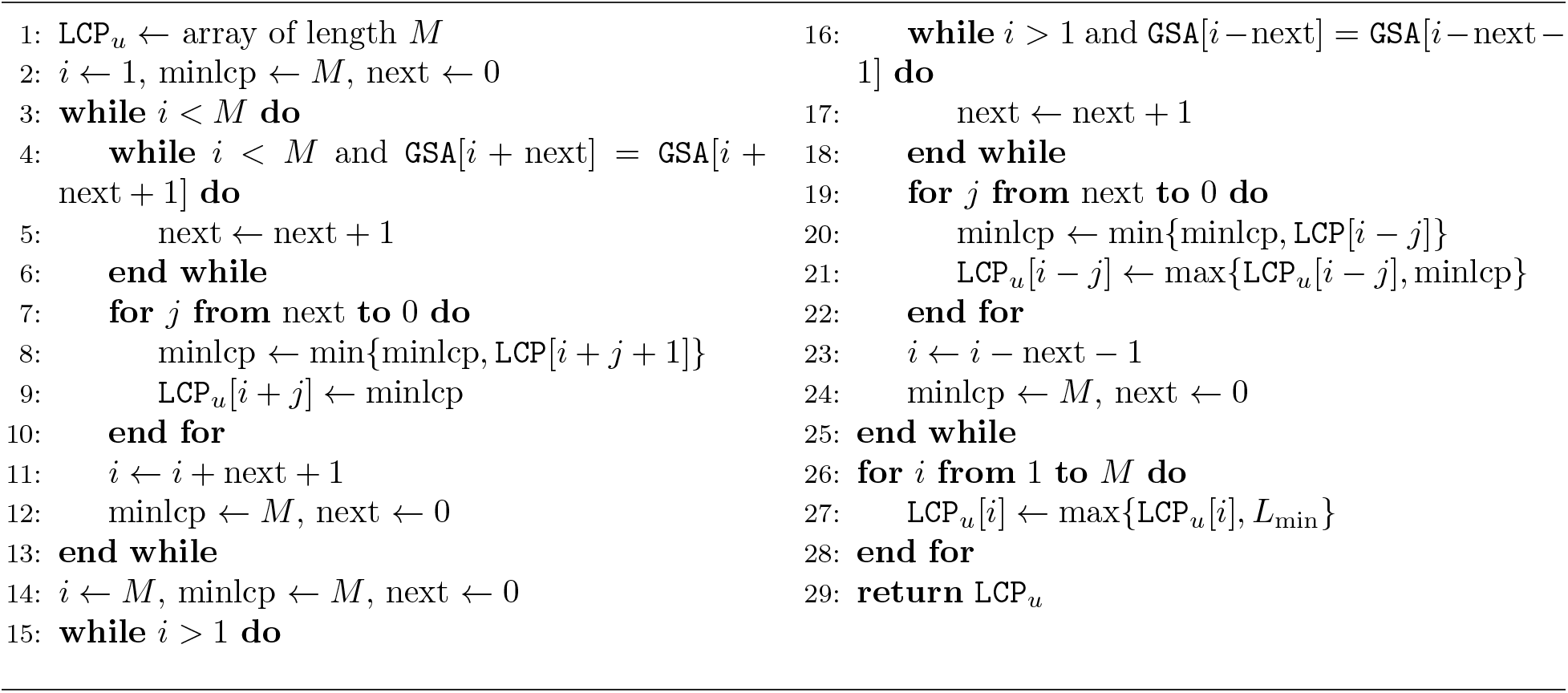
LCP_*u*_FromLCP(`GSA`, `LCP`, *L*_min_)

#### Lemma 4.

*Both LCP_*u*_FromLCP and LCP*_*d*_*FromLCP run in O(M) time*.

*Proof.* Through a simple aggregate analysis, we can see that LCP_*u*_FromLCP visits each entry of `GSA`, `LCP` and `LCP`_*u*_ 2 times; LCP_*d*_FromLCP* visits each entry of `GSA`, `LCP` 3 times and each entry of 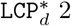 2 times for either focus = +/−.

Combining the above lemmata, we concluse that

#### Theorem 5.

*Both `SU` and `SD` can be computed in in O(M) time*.

### 5.3 Sampling unique substrings

Recall that 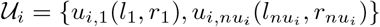 defines either the collection of all shortest unique substrings or unique substrings with minimum length *L*_min_ on a given genome *s*_*i*_ (sorted by *l*_*x*_, namely 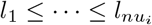). In fact, the list of left indices 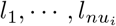 are stored in the corresponding 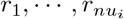 entries in `SU` array. Due to the minimum length constraint, no substring 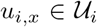 can be a substring of any other 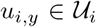 if they are not identical (in fact, there could be some 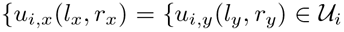 for *x* ≠ *y*). This makes 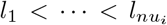 and 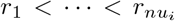. The goal of sampling unique substrings from 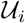 is to identify and maintain the smallest number of unique substrings such that they cover the same set of unique *L*-mers on *s*_*i*_ as 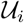 (if *L*_max_ = *L* then the sampled unique substrings should cover all unique *L*-mers).

Here we present the greedy sampling strategy implemented by `CAMMiQ` to sample unique substrings from 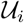. Denote by begin_*i*_ the beginning position of *s*_*i*_ in *s* and by 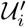 the unique substrings already sampled (initially 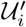 is empty). Starting with begin_*i*_, consider every *L*-mer of *s*_*i*_ from left to right; if it does not include any unique substring, then ignore this *L*-mer; otherwise add its rightmost unique substring into 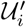 and move to the next *L*-mer which does not include this substring until reaching the *L*-mer that ends at begin_*i*_ + |*s*_*i*_| − 1. At the and of this, add the sampled unique substrings in 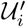 to the hash table described in Section 2.1.

**Algorithm 3.**
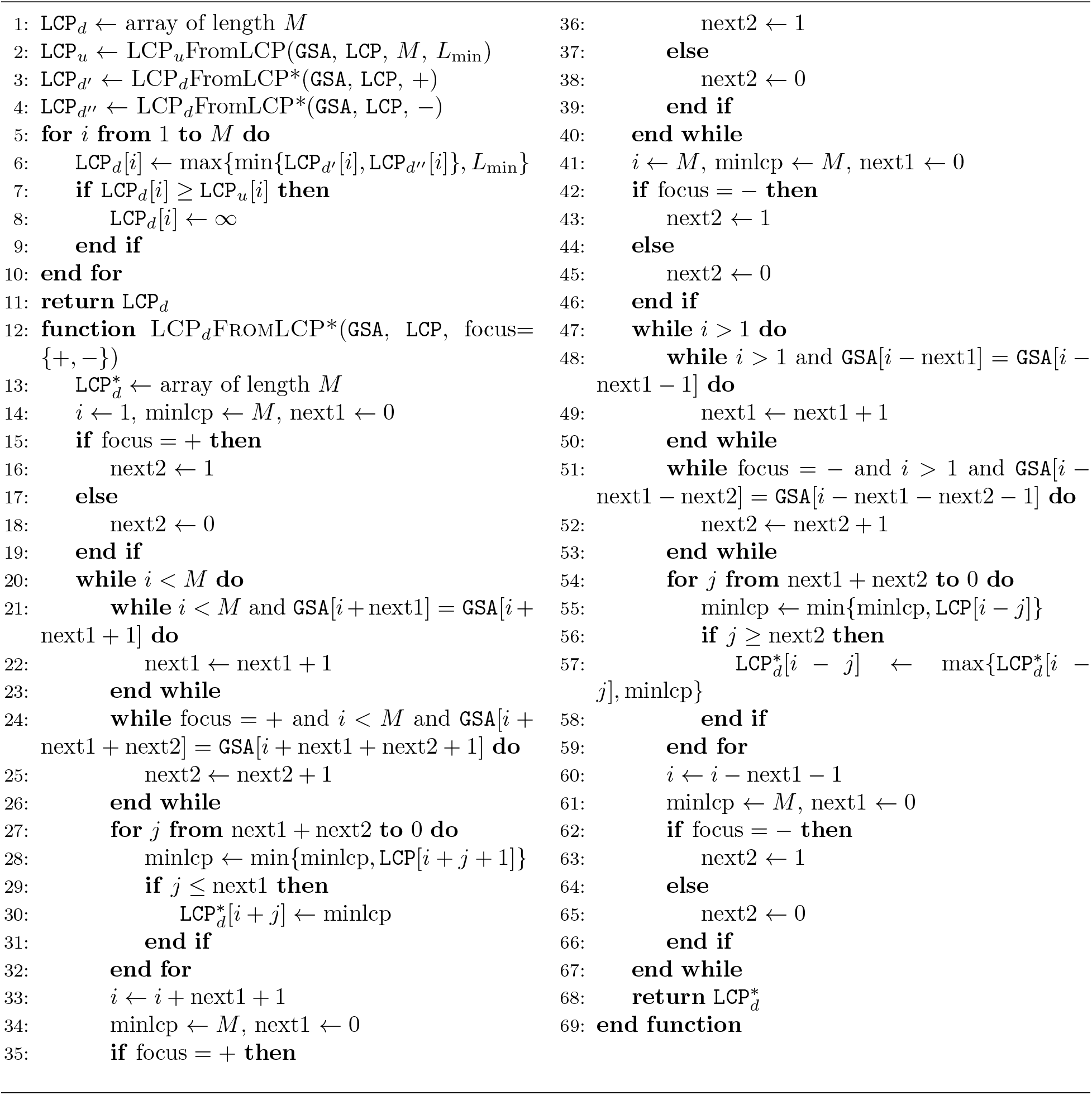
LCP_*d*_FromLCP(`GSA`, `LCP`, *L*_min_)

In the following we show that the above greedy strategy obtains the smallest number of unique substrings that cover the same set of unique *L*-mers as 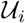, provided that each unique substring in 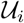 occurs only one time (i.e., any 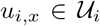 is not identical to another 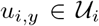 if *x* ≠ *y*). As a result, the total number of unique substrings in 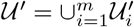 included in `CAMMiQ` index is also as small as possible.

#### Theorem 6.

*If* 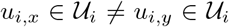 *for x ≠ y, then GreedySampling returns the smallest* 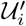 *such that if an L-mer includes some* 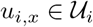, *then it also includes at least one* 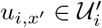.

*Proof.* Consider 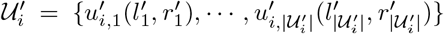 that GreedySampling returns; also consider an alternative sample 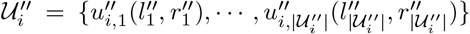 of 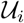 that covers the same set of *L*-mers; assume both sets are sorted by the left indices 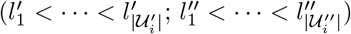. First, observe that 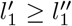 since 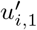 is the rightmost unique substring in 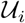 which is fully included in the leftmost unique *L*-mer. As a consequence we must have 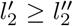 (and so on). Otherwise, if 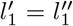 then 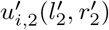 is not the rightmost unique substring in the next unique *L*-mer whose left index is greater than 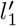 and if 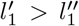 then there is some unique *L*-mer not covered by any unique substring in 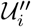. Therefore 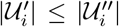 since there is a injection between the elements in 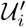 and 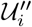 until reaching the last unique *L*-mer on *s*_*i*_.

**Algorithm 4.**
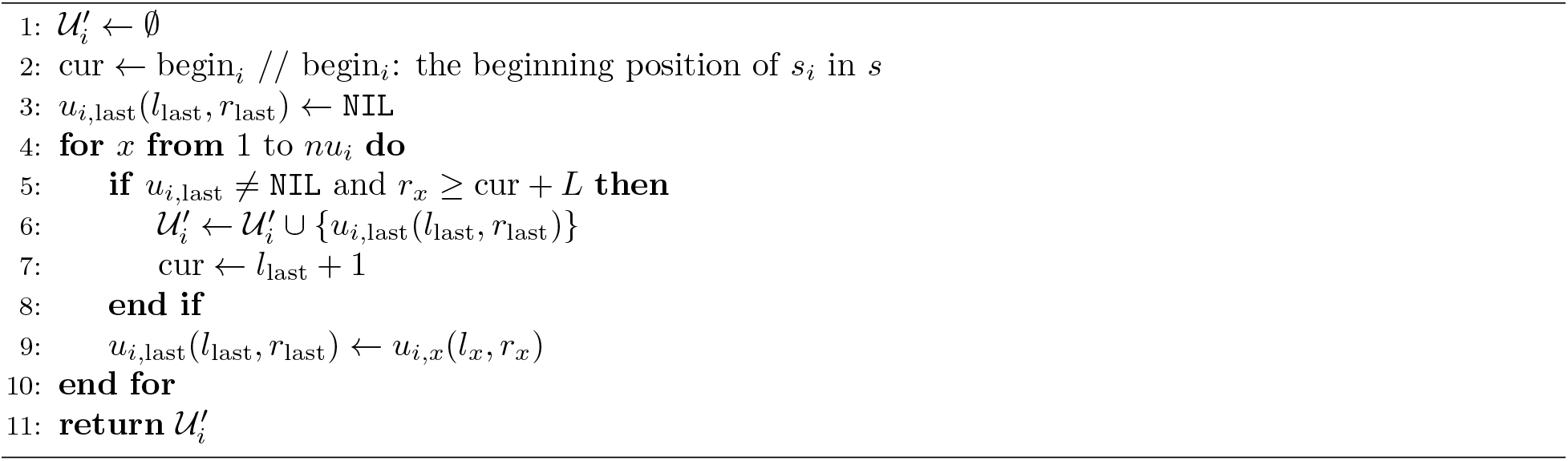
GreedySampling(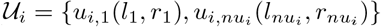, *L*)

#### Corollary 7.

*If* 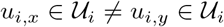 *for x ≠ y on each s*_*i*_, *then* 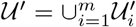 *is the smallest set of unique substrings such that if an L-mer on s*_*i*_ *includes some* 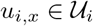, *then it also includes at least one* 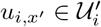.

We note that the greedy sampling strategy works in practice even if there are actually unique substrings occurring more than once in a given genome, meaning 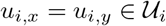 for some *y* > *x*, leading to 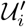 (and thus 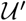) being close to optimality, since these unique substrings would constitute a very small proportion (≤ 0.1%) with the default minimum length *L*_min_ = 26 of unique substrings. This gives significantly smaller indices than alternative *k*-mer based tools, and results in an integer program with a small number of constraints.

We applied the above strategy to sample doubly-unique substrings in 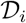, to obtain the minimum size 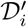 for each genome *s*_*i*_ so that the aggregate set of doubly-unique substrings 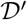 is at most twice the optimal, provided that each doubly-unique substring occurs once in each of the corresponding genomes.

#### Corollary 8.

*If* 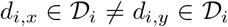 *for x ≠ y on each s_i_*, *then* 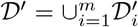 *is at most twice as large as the smallest set of doubly-unique substrings such that if an L-mer on s_i_ includes some* 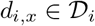, *then it also includes at least one* 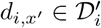.

### 5.4 Proof of Theorem 1

We begin with the following theorem from Weissman et al. [62]. that bounds the L1 distance between the empirical distribution of a sequence of independent, identically distributed random variables and the true distribution.

#### Theorem 9.

*Let P be a probability distribution on the set* 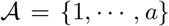. *Let X_1_,X_2_, ···, X_n_ be i.i.d. random variables distributed according to P. Then, for any given ϵ* > 0,

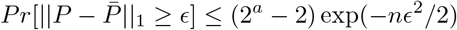

*where* 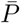 *is the empirical estimation of P defined as* 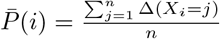, *where Δ(e)* = 1 *if and only if e is true and* Δ(*e*) = 0 *otherwise*.

Now consider a collection of genomes 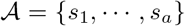 with relative abundances *p*_1_, ···, *p*_*a*_ and the set 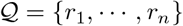 of *n* reads (i.e., *L*-mers) sampled independently and uniformly at random from 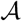 according to *p*_1_, ···, *p*_*a*_. On each genome *s*_*i*_ let 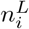 denote the total number of *L*-mers and 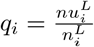 be the proportion of unique *L*-mers; then the probability of a read 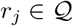 corresponds to a unique *L*-mer on *s*_*i*_ is 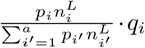, and the probability of a read 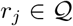 does not correspond to any unique *L*-mers on *s*_*i*_ is 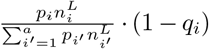.

Therefore *r*_1_, ···, *r*_*j*_ are i.i.d. distributed according to 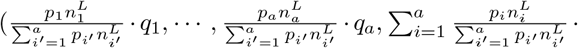. 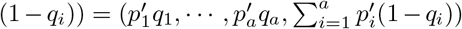 where the last term corresponds to the reads that are not unique to any 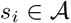.

**Proof of Theorem 1.** Let *c*_*i*_ be the number of reads assigned to *s*_*i*_. Also let 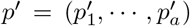. We set 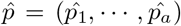, by defining 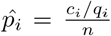 to be the predicted abundance of *s*_*i*_ based on the number of reads assigned to it. Consider 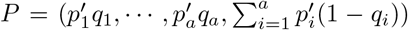 and 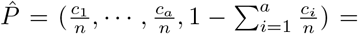 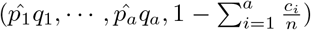; by definition, we have 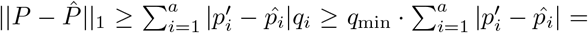 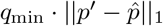. Then the following three theorem statements hold.

- (i) Given that 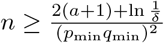, we have

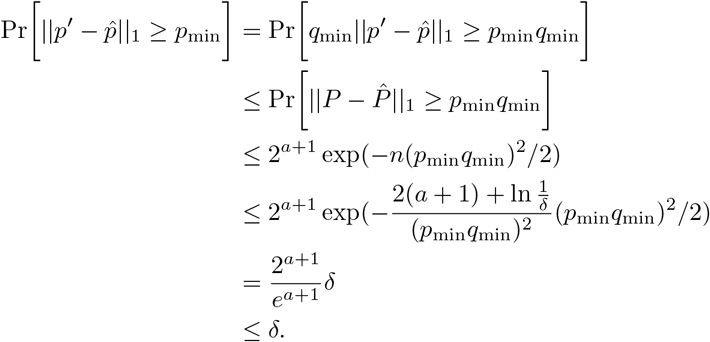 This implies that with probability ≥ 1 − *δ* the L1 distance between *p′* and 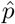 is upper bounded by *p*_min_. As a result we have 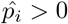 for each 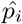, i.e. *c*_*i*_ ≥ 0.
- (ii) The proof follows by simply replacing *p*_min_ with *ϵ* in the proof of (i).
- The proof follows by simply replacing *p*_min_ with 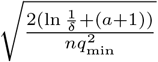 in the proof of (i). Specifically,

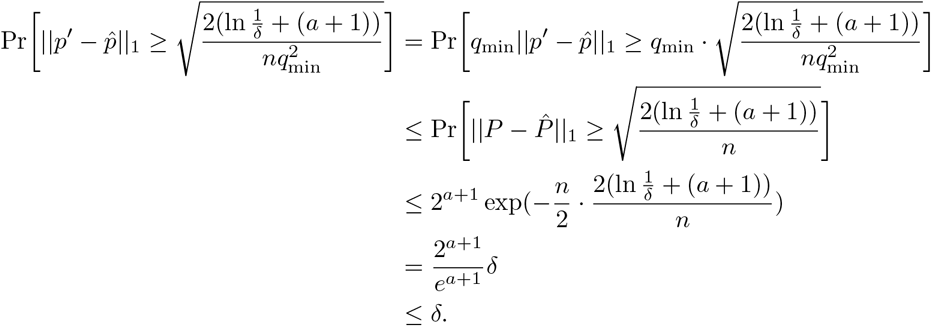

### 5.5 blastn analysis of *Salmonella* strains

The RNASeq reads for each cell studied in 58. are stored as separate read sets in the Sequence Read Archive (SRA) [63]. To reduce the number of reads that need to be downloaded or aligned, we used ReadFinder (https://github.com/morgulis/ReadFinder) to find any reads that could plausibly map to either the *Salmonella* strains *LT2* or *D23580* with permissive parameters allowing alignments that stray by as much as four diagonals from the main diagonal. ReadFinder uses similar methods as SRPRISM 64. to search SRA without the need to download the data and is more streamlined than SRPRISM for our purpose of finding candidate matching reads. We then used blastn 65. with word size 16 to find local (and ideally, global) alignments between the reads identified by ReadFinder and either *Salmonella* strain. This test is much simpler than the `CAMMiQ` test because we did not consider alignments to any *Salmonella* strains other than the two actually used in the original wet lab experiment.

Most reads that align to either strain actually align to the two strains equally well. To decide which reads align strictly better to one strain or the other, we implemented an in-house script with the following rules. A blastn alignment is “high-quality” if it has length at least 70 (taking into account that the reads are of length 75), has identity percentage ≥ 98. A read *R* maps better to the *LT2* strain if either:

- *R* has a high-quality alignment to the *LT2* strain but R has no high-quality alignment to the *D23580* strain or
- *R* has high-quality alignments to both strains and the best *LT2* alignment is at least as good as the best *D23580* alignment on i) length ii) identity percentage, iii) gaps and is better than the best D-strain alignment on at least one of the thee Roman numeral criteria.

The criteria for mapping better to the D-strain are symmetric. The script reports counts of how many reads map align strictly better to each of the two strains. Although the data in [58]. consist of paired reads, each mate pair was treated individually in the blastn analysis.

## 6 Supplementary Figures

Figure 5 presents 50 genomes in our species-level dataset with the least number of unique *L*-mers. In this graph each node represents one such genome; each edge connects two nodes if they share a doubly-unique substring. Solid black edges indicate a pair of nodes that share at least 30 doubly-unique substrings; the remaining edges in grey indicate node pairs with fewer number of shared doubly-unique substrings. Notice that there is a special node in the center, representing the union of all genomes not included in these 50-genome subset. Any node connected to this special node by a single edge, or by a path, is identifiable and quantifiable by `CAMMiQ`, provided that all edges in this path are black (22 of these 50 nodes are as such) or all nodes in this path have sufficient abundance. Note that 20 of the genomes here are connected to this special node by a black edge: these are the genomes that form the least-quantifiable-20 dataset in our experiments.

**Figure 5:**
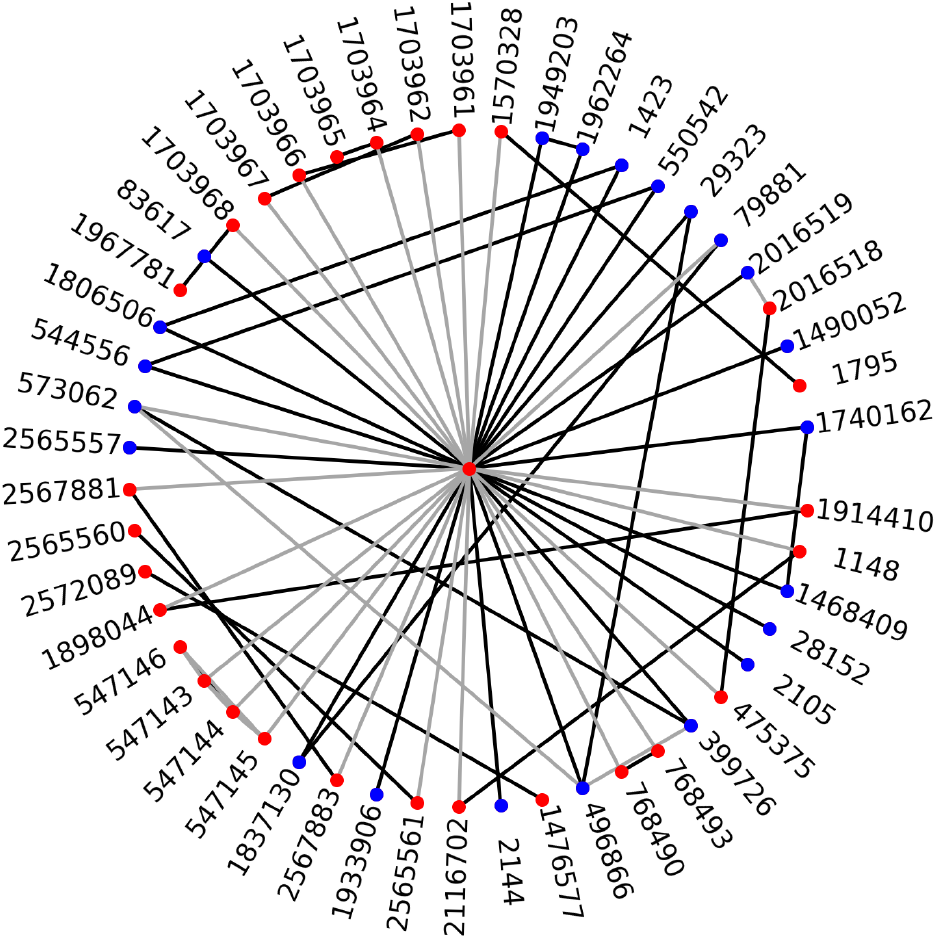
Shared doubly-unique substrings among the 50 genomes with the least number of unique *L*-mers in our species-level dataset consisting of 4122 RefSeq bacteria genomes. Each node represents one of these 50 genomes, labeled with its NCBI taxID at species level. The central node specially represents the remaining 4072 genomes. An edge connecting two nodes indicate at least one doubly-unique substring shared between them. A black edge indicates ≥ 30 doubly-unique substrings in `CAMMiQ`’s index shared between the two corresponding genomes. All other edges in grey imply < 30 shared doubly-unique substrings. A blue-colored node indicates one that is connected to the central node through a path of black edges. As such, they are relatively easy to identify and quantify; 22 of these 50 nodes are blue. The remaining (red) nodes can be identified by `CAMMiQ` provided they have “sufficient abundance” in the query.

## Footnotes

1 There are also the so-called “gapped” *k*-mers used in a variety of applications such as functional classification of metagenomic reads [49]; however not only these applications are different from that of `CAMMiQ`, but also, they use fixed length substrings. Finally, there are alternative approaches using, e.g. linked-reads [50]; `CAMMiQ` focuses on standard Illumina reads to be able to analyze a wealth of readily available sequencing data sets.

2 We explain how `CAMMiQ` resolves ambiguity on reads that include substrings which suggest conicting assignments later in the paper.

3 The complete set of genomes in this database is 617 but only 614 can be downloaded from RefSeq.

4 This happens if the read includes one or more doubly unique substrings from the same pair of genomes but no unique substring.

5 Here we used the latest set of marker genes mpa v20 m200 in MetaPhlAn2.

6 As can be seen in Figure 4B, even though CAMMiQ’s 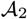 type queries generally improve over its 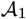 type queries with respect to the number of cells correctly identified with STM-LT2 strain or STM-D23580 strains, it seems to also introduce some potential false positive calls (e.g. the last panel in the figure corresponding to the controls). This could be due to additional reads utilized by 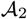 queries impacted by read errors or misassigned reads.

## References

[1] Huttenhower, C. et al. Structure, function and diversity of the healthy human microbiome. nature 486, 207 (2012).

[2] Bullman, S. et al. Analysis of fusobacterium persistence and antibiotic response in colorectal cancer. Science 358, 1443–1448 (2017).

[3] Castellarin, M. et al. Fusobacterium nucleatum infection is prevalent in human colorectal carcinoma. Genome Research 22, 299–306 (2012).

[4] Gur, C. et al. Binding of the fap2 protein of fusobacterium nucleatum to human inhibitory receptor tigit protects tumors from immune cell attack. Immunity 42, 344–355 (2015).

[5] Gur, C. et al. Fusobacterium nucleatum supresses anti-tumor immunity by activating ceacam1. Oncoimmunology 8, e1581531 (2019).

[6] Kostic, A. D. et al. Genomic analysis identifies association of fusobacterium with colorectal carcinoma. Genome Research 22, 292–298 (2012).

[7] Yu, T. et al. Fusobacterium nucleatum promotes chemoresistance to colorectal cancer by modulating autophagy. Cell 170, 548–563 (2017).

[8] Huson, D. H., Auch, A. F., Qi, J. & Schuster, S. C. Megan analysis of metagenomic data. Genome Research 17, 377–386 (2007).

[9] Altschul, S. F., Gish, W., Miller, W., Myers, E. W. & Lipman, D. J. Basic local alignment search tool. Journal of Molecular Biology 215, 403–410 (1990).

[10] Walker, M. A. et al. Gatk pathseq: a customizable computational tool for the discovery and identification of microbial sequences in libraries from eukaryotic hosts. Bioinformatics 34, 4287–4289 (2018).

[11] Poore, G. D. et al. Microbiome analyses of blood and tissues suggest cancer diagnostic approach. Nature 579, 567–574 (2020).

[12] Wood, D. E. & Salzberg, S. L. Kraken: ultrafast metagenomic sequence classification using exact alignments. Genome Biology 15, R46 (2014).

[13] Elworth, R. et al. To petabytes and beyond: recent advances in probabilistic and signal processing algorithms and their application to metagenomics. Nucleic Acids Research (2020).

[14] Robinson, W., Schischlik, F., Gertz, E. M., Schäffer, A. A. & Ruppin, E. Identifying the landscape of intratumoral microbes via a single cell transcriptomic analysis. bioRxiv (2020).

[15] Liu, B., Gibbons, T., Ghodsi, M., Treangen, T. & Pop, M. Accurate and fast estimation of taxonomic profiles from metagenomic shotgun sequences. Genome Biology 12 (Suppl 2), S4 (2011).

[16] Segata, N. et al. Metagenomic microbial community profiling using unique clade-specific marker genes. Nature Methods 9, 811 (2012).

[17] Lopresti, D. & Tomkins, A. Block edit models for approximate string matching. Theoretical Computer Science 181, 159–179 (1997).

[18] Cormode, G., Paterson, M., Sahinalp, S. C. & Vishkin, U. Communication complexity of document exchange. In Proceedings of the eleventh annual ACM-SIAM symposium on Discrete algorithms, 197–206 (Society for Industrial and Applied Mathematics, 2000).

[19] Li, M., Chen, X., Li, X., Ma, B. & Vitányi, P. M. The similarity metric. IEEE Transactions on Information Theory 50, 3250–3264 (2004).

[20] Vinga, S. & Almeida, J. Alignment-free sequence comparison—a review. Bioinformatics 19, 513–523 (2003).

[21] Leimeister, C.-A. & Morgenstern, B. Kmacs: the k-mismatch average common substring approach to alignment-free sequence comparison. Bioinformatics 30, 2000–2008 (2014).

[22] Ames, S. K. et al. Scalable metagenomic taxonomy classification using a reference genome database. Bioinformatics 29, 2253–2260 (2013).

[23] Břinda, K., Sykulski, M. & Kucherov, G. Spaced seeds improve k-mer-based metagenomic classification. Bioinformatics 31, 3584–3592 (2015).

[24] Kawulok, J. & Deorowicz, S. Cometa: classification of metagenomes using k-mers. PloS ONE 10, e0121453 (2015).

[25] Tu, Q., He, Z. & Zhou, J. Strain/species identification in metagenomes using genome-specific markers. Nucleic Acids Research 42, e67–e67 (2014).

[26] Ounit, R., Wanamaker, S., Close, T. J. & Lonardi, S. Clark: fast and accurate classification of metage-nomic and genomic sequences using discriminative k-mers. BMC Genomics 16, 236 (2015).

[27] Luo, Y., Zeng, J., Berger, B. & Peng, J. Low-density locality-sensitive hashing boosts metagenomic binning. In International Conference on Research in Computational Molecular Biology, 255 (Springer, 2016).

[28] Ondov, B. D. et al. Mash: fast genome and metagenome distance estimation using minhash. Genome Biology 17, 132 (2016).

[29] McHardy, A. C., Martín, H. G., Tsirigos, A., Hugenholtz, P. & Rigoutsos, I. Accurate phylogenetic classification of variable-length dna fragments. Nature Methods 4, 63 (2007).

[30] Rosen, G., Garbarine, E., Caseiro, D., Polikar, R. & Sokhansanj, B. Metagenome fragment classification using n-mer frequency profiles. Advances in Bioinformatics 2008 (2008).

[31] Brady, A. & Salzberg, S. L. Phymm and phymmbl: metagenomic phylogenetic classification with interpolated markov models. Nature Methods 6, 673 (2009).

[32] Rosen, G. L., Reichenberger, E. R. & Rosenfeld, A. M. Nbc: the naive bayes classification tool webserver for taxonomic classification of metagenomic reads. Bioinformatics 27, 127–129 (2010).

[33] Vervier, K., Mahé, P., Tournoud, M., Veyrieras, J.-B. & Vert, J.-P. Large-scale machine learning for metagenomics sequence classification. Bioinformatics 32, 1023–1032 (2015).

[34] Anyansi, C., Straub, T. J., Manson, A. L., Earl, A. M. & Abeel, T. Computational methods for strain-level microbial detection in colony and metagenome sequencing data. Frontiers in Microbiology 11, 1925 (2020).

[35] Marshall, J. A. Mixed infections of intestinal viruses and bacteria in humans. In Polymicrobial Diseases (ASM Press, 2002).

[36] Cohen, T. et al. Mixed-strain mycobacterium tuberculosis infections and the implications for tubercu-losis treatment and control. Clinical microbiology reviews 25, 708–719 (2012).

[37] Balmer, O. & Tanner, M. Prevalence and implications of multiple-strain infections. The Lancet infectious diseases 11, 868–878 (2011).

[38] Secher, T., Brehin, C. & Oswald, E. Early settlers: which e. coli strains do you not want at birth? American Journal of Physiology: Gastrointestinal and Liver Physiology 311, G123–G129 (2016).

[39] Gerner-Smidt, P. et al. Whole genome sequencing: Bridging one-health surveillance of fooborne diseases. Frontiers in Public Health 7, 172 (2019).

[40] Lin, Y.-Y. et al. Cliiq: Accurate comparative detection and quantification of expressed isoforms in a population. In International Workshop on Algorithms in Bioinformatics, 178–189 (Springer, 2012).

[41] Li, W., Feng, J. & Jiang, T. Isolasso: a lasso regression approach to rna-seq based transcriptome assembly. Journal of Computational Biology 18, 1693–1707 (2011).

[42] Dao, P. et al. Orman: optimal resolution of ambiguous rna-seq multimappings in the presence of novel isoforms. Bioinformatics 30, 644–651 (2014).

[43] Sobih, A., Tomescu, A. I. & Mäkinen, V. Metaflow: Metagenomic profiling based on whole-genome coverage analysis with min-cost flows. In International Conference on Research in Computational Molecular Biology, 111–121 (Springer, 2016).

[44] Breitwieser, F., Baker, D. & Salzberg, S. L. Krakenuniq: confident and fast metagenomics classification using unique k-mer counts. Genome Biology 19, 1–10 (2018).

[45] Solomon, B. & Kingsford, C. Fast search of thousands of short-read sequencing experiments. Nature Biotechnology 34, 300 (2016).

[46] Solomon, B. & Kingsford, C. Improved search of large transcriptomic sequencing databases using split sequence bloom trees. In International Conference on Research in Computational Molecular Biology, 257–271 (Springer, 2017).

[47] Sun, C., Harris, R. S., Chikhi, R. & Medvedev, P. Allsome sequence bloom trees. In International Conference on Research in Computational Molecular Biology, 272–286 (Springer, 2017).

[48] Ferdman, M., Johnson, R. & Patro, R. Mantis: A fast, small, and exact large-scale sequence-search index. In Research in Computational Molecular Biology, 271 (Springer, 2018).

[49] Nazeen, S., Yu, Y. W. & Berger, B. Carnelian uncovers hidden functional patterns across diverse study populations from whole metagenome sequencing reads. Genome biology 21, 1–18 (2020).

[50] Danko, D. C., Meleshko, D., Bezdan, D., Mason, C. & Hajirasouliha, I. Minerva: an alignment-and reference-free approach to deconvolve linked-reads for metagenomics. Genome research 29, 116–124 (2019).

[51] Matias, Y., Muthukrishnan, S., Sahinalp, S. C. & Ziv, J. Augmenting suffix trees, with applications. In European Symposium on Algorithms, 67–78 (Springer, 1998).

[52] Kasai, T., Lee, G., Arimura, H., Arikawa, S. & Park, K. Linear-time longest-common-prefix computation in suffix arrays and its applications. In Annual Symposium on Combinatorial Pattern Matching, 181–192 (Springer, 2001).

[53] Kärkkäinen, J. & Sanders, P. Simple linear work suffix array construction. In International Colloquium on Automata, Languages, and Programming, 943–955 (Springer, 2003).

[54] Ilie, L. & Smyth, W. F. Minimum unique substrings and maximum repeats. Fundamenta Informaticae 110, 183–195 (2011).

[55] Karp, R. M. & Rabin, M. O. Efficient randomized pattern-matching algorithms. IBM journal of research and development 31, 249–260 (1987).

[56] Vazirani, V. V. Approximation algorithms (Springer Science & Business Media, 2013).

[57] Pruitt, K. D., Tatusova, T. & Maglott, D. R. Ncbi reference sequences (refseq): a curated non-redundant sequence database of genomes, transcripts and proteins. Nucleic Acids Research 35, D61–D65 (2007).

[58] Aulicino, A. et al. Invasive salmonella exploits divergent immune evasion strategies in infected and bystander dendritic cell subsets. Nature Communications 9, 4883 (2018).

[59] Wood, D. E., Lu, J. & Langmead, B. Improved metagenomic analysis with kraken 2. Genome Biology 20, 257 (2019).

[60] Truong, D. T. et al. Metaphlan2 for enhanced metagenomic taxonomic profiling. Nature Methods 12, 902 (2015).

[61] Forster, S. C. et al. A human gut bacterial genome and culture collection for improved metagenomic analyses. Nature Biotechnology 37, 186 (2019).

[62] Weissman, T., Ordentlich, E., Seroussi, G., Verdu, S. & Weinberger, M. J. Inequalities for the L1 deviation of the empirical distribution. Hewlett-Packard Labs, Tech.Rep (2003).

[63] Leinonen, R., Sugawara, H., Shumway, M. et al. The sequence read archive. Nucleic Acids Research 39, D19–D21 (2011).

[64] Morgulis, A. & Agarwala, R. Srprism (single read paired read indel substitution minimizer): an efficient aligner for assemblies with explicit guarantees. GigaScience 9, giaa023 (2020).

[65] Altschul, S. F. et al. Gapped blast and psi-blast: a new generation of protein database search programs. Nucleic Acids Research 25, 3389–3402 (1997).

